# The Effect of Alterations of Schizophrenia-Associated Genes on Gamma Band Oscillations

**DOI:** 10.1101/2020.09.28.316737

**Authors:** Christoph Metzner, Tuomo Mäki-Marttunen, Gili Karni, Hana McMahon-Cole, Volker Steuber

## Abstract

Abnormalities in the synchronized oscillatory activity of neurons in general and, specifically in the gamma band, might play a crucial role in the pathophysiology of schizophrenia. While these changes in oscillatory activity have traditionally been linked to alterations at the synaptic level, we demonstrate here, using computational modeling, that common genetic variants of ion channels can contribute strongly to this effect. Our model of primary auditory cortex highlights multiple schizophrenia-associated genetic variants that reduce gamma power in an auditory steady-state response task. Furthermore, we show that combinations of several of these schizophrenia-associated variants can produce similar effects as the more traditionally considered synaptic changes. Overall, our study provides a mechanistic link between schizophrenia-associated common genetic variants, as identified by genome-wide association studies, and one of the most robust neurophysiological endophenotypes of schizophrenia.

## 1 Introduction

The search for biological causes of psychiatric disorders has up to now met with limited success. While genetics and basic neuroscience have both made tremendous advances over the last decade, mechanistic links between genetic findings and clinical symptoms have so far not been discovered. Many have argued that symptom-based classifications of psychiatric illnesses might not be possible to map to alterations at the microscopic scale [18, 45], and have proposed to use biomarkers or endophenotypes, which in turn might correlate more clearly with genetic variants [45]. For example, the biomarkers and endophenotypes of schizophrenia (SCZ) include reduced mismatch negativity [100], reduced pre-pulse inhibition [96] and changes to evoked and induced oscillations in multiple frequency bands in a large variety of tasks (e.g. [99]). Importantly, recent advances in computational modelling allow for the integration of knowledge about genetic contributions to ion channels and excitability and can be used to predict changes to macroscopic electroencephalography (EEG) or magnetoencephalography (MEG) signals ([35, 107, 60], for a review of this emerging subfield of computational psychiatry see [62]). For example, simulations of a detailed model of tufted layer 5 pyramidal cells have recently been used to predict the effect of SCZ-associated variants of ion channel-encoding genes on neural activity in the delta frequency band [63].

In general, oscillations in the low and high frequency range enable coordinated interactions between distributed neuronal responses [9, 27, 8, 87] and have been demonstrated to be functionally relevant [97]. For example, gamma oscillations have been linked to perception [34], attention [28], memory [92], consciousness [67] and synaptic plasticity [109]. In patients with schizophrenia, gamma power and coherence have been consistently found to be decreased during neural entrainment [51, 104, 49, 95] as well as during several sensory (e.g. visual Gestalt [90]) and cognitive (e.g. working memory [15]) tasks. Importantly, schizophrenia is also associated with disturbances in many of the above mentioned functions [77, 29, 98]. Mathematical analyses and computer simulations have demonstrated that gamma oscillations arise through the local interplay between excitatory and inhibitory populations, either through tonic excitation of inhibitory cells and subsequent rhythmic inhibition of excitatory cells (interneuron gamma or ING) or through rhythmic excitation of inhibitory cells and subsequent rhythmic inhibition of excitatory cells (pyramidal-interneuron gamma or PING) [111, 6]. The anatomical and electrophysiological properties of a particular subtype of inhibitory interneurons, the parvalbumin-positive (PV^+^) interneurons, make them ideally suited for the fast, strong and temporally precise inhibition necessary for the generation of gamma rhythms [32]. Furthermore, optogenetically driving PV^+^ interneurons was found to enhance gamma rhythms [11]. Consequently, cellular level alterations at PV^+^ interneurons in schizophrenia have been linked to well-known auditory steady-state response (ASSR) deficits in the gamma band [104, 70, 71, 47, 86]. However, these studies have focused on changes to the strength and temporal dynamics of synaptic transmission. While synaptic transmission dynamics undoubtedly play a crucial role in the generation of neural oscillations, cell-intrinsic properties such as ionic conductances can also alter the ability of a network to generate and maintain oscillations. This is of particular importance since many of the recently discovered gene variants associated with schizophrenia relate to ionic channels or Ca^2+^ transporters [83, 22].

In this study, we use an established framework to translate the effect of common single-nucleotide polymorphism (SNP) variants associated with schizophrenia into a biophysically detailed model of a layer 5 pyramidal cell [60]. We then use a morphologically reduced version of the cell [59] together with a model of a PV^+^ interneuron [103] in a microcircuit model and explore the effect of the genetic variants on gamma entrainment. We demonstrate that while single gene variants typically only have small effects on gamma auditory steady-state entrainment, combinations of them can reduce the entrainment comparable to the synaptic alterations mentioned above and replicate observations in schizophrenia patients. Our findings therefore provide a mechanistic link between the scale of single genes and an important endophenotype of schizophrenia. Furthermore, the proposed model represents an ideal test-bed for the identification of targets for potential pharmacological agents aiming to reverse gamma deficits in schizophrenia.

## 2 Methods

### 2.1 Network Model

This study was based on a high-complexity, biophysically detailed model of thick-tufted layer 5 pyramidal cells [37]. The model includes a detailed reconstructed morphology, models of the dynamics of eleven different ionic channels and a description of the intracellular Ca^2+^ concentration [37]. Following earlier work, we incorporated human *in vitro* electrophysiological data on ion channel behavior from the functional genomics literature into this model [60, 63]. Due to the computational complexity of the original model, consisting of 196 compartments, we decided to use a reduced-morphology model, where passive parameters, ion channel conductances and parameters describing Ca^2+^ dynamics were fitted to reproduce the behaviour of the original model [59]. The inhibitory cells in the network were based on a model of fast-spiking PV^+^ basket cells taken from [103]. These two single cell models were combined into a microcircuit network model consisting of 256 excitatory and 64 inhibitory neurons. Cells were connected via AMPA and NMDA receptor-mediated synaptic currents in the case of excitatory connections and GABA_*A*_ receptor-mediated synaptic currents for inhibitory connections. Additionally, model cells received two types of input, Poissonian noise to all cells representing background activity in the cortex and rhythmic input representing the sensory input during auditory entrainment. Note that a smaller percentage of inhibitory interneurons (35%) received no sensory input drive; this reflects preferential thalamic drive to pyramidal cell populations [4], which has also been used in other models [104]. This was to ensure that a subpopulation of the inhibitory neurons had a weak enough drive to be dominated by pyramidal cell activity. This subpopulation was necessary to maintain a 20 Hz peak for 40 Hz drive in of the synaptic alteration conditions against which we compared the genetic alterations. For a detailed discussion see the study by Vierling-Claassen et al. [104].

### 2.2 Integration of Genetic Variants

We closely follow earlier work of [60, 64, 63] to integrate the effect of SNP-like genetic variants into our network model (details in Supplementary Section 4). In summary, we selected a set of genes, restricted to ion-channel-encoding genes likely to be expressed in layer 5 pyramidal cells, obtained from a large genome-wide association study (GWAS) [83]. For this set of genes, we searched the literature for genetic variants and their effects on electrophysiological parameters of pyramidal cells. This left us with 86 variants of the following genes: CACNA1C, CACNA1D, CACNB2, SCN1A and, HCN1 [50, 19, 40, 91, 94, 93, 5, 112, 79, 80, 3, 54, 17, 66, 55, 43, 14, 102, 105, 13, 65, 44, 53, 108]. These variants typically had a large effect on the electrophysiology of layer 5 pyramidal cells. However, due to the polygenic nature of SCZ, it can be assumed that the risk of the disorder is not caused by a single SCZ-associated SNP and, therefore, one would not expect large effects from the common variants identified in GWAS studies. Subsequently, we applied a downscaling procedure, as outlined in [60, 63]. In short, we downscaled the changes of model parameters induced by a variant (multiplication by a factor either on a linear or logarithmic scale, depending on the type of the parameter) until the cell response to predefined stimuli stayed within a certain range (Supplementary Section 4). This resulted in a set of 86 ‘small-effect’ model variants, which were used as models for the effects of common variants on layer 5 pyramidal cell electrophysiological response features. As in earlier studies using this approach [60, 63], we will use the term ‘variant’ for a genetic variant in a human or animal genome and the term ‘model variant’ for a model of a gene variant constructed as described above.

## 3 Results

We executed simulations of the network model with background synaptic noise and a periodic drive at 40 Hz, mimicking auditory steady-state stimulation experiments. For each model variant in Supplementary table S1, we repeated the set of 200 simulations (10 ‘trials’ for each of the 20 ‘virtual subjects’, see Supplementary section 4). For each of these model variants, the parameters of the ion-channels were altered in a subtle way, leading to changes in their activation, and subsequently to modified network dynamics and gamma entrainment.

### 3.1 Single Variants Can Produce ASSR Power Deficits in the Gamma Range

First, we analyzed the evoked ASSR power in the gamma range. Most of the model variants altered gamma power in a weak or moderate way and the model variants could both increase or decrease gamma power (Figure 2). Model variants affecting the *Na*^+^ channels had hardly any effect on entrainment power which reflects the small scaling coefficients imposed by the downscaling scheme (see Supplementary Table 3). Four model variants, one affecting the CACNA1C gene, one affecting the CACNA1I gene and two affecting the HCN1A gene, led to strong decreases of evoked power (Figure 3). For the model variants affecting *I*_*CaHV A*_, the change in evoked gamma power was positively correlated with the offset of the activation variable *m* (Pearson correlation: *r* = 0.53, *p* < 0.001; Supplementary Figure S9) but there was no significant correlation with other parameters describing the ion channel dynamics (Supplementary Figure S9). Furthermore, no model variant strongly increased the 20 Hz component (see Supplementary Figure S8), i.e. we observed no shift to the first subharmonic of the 40 Hz drive, which would be indicative of a ‘beat-skipping’ behaviour, where the inhibition suppresses every other drive stimulus as seen in models of altered synaptic dynamics [104].

**Figure 1:**
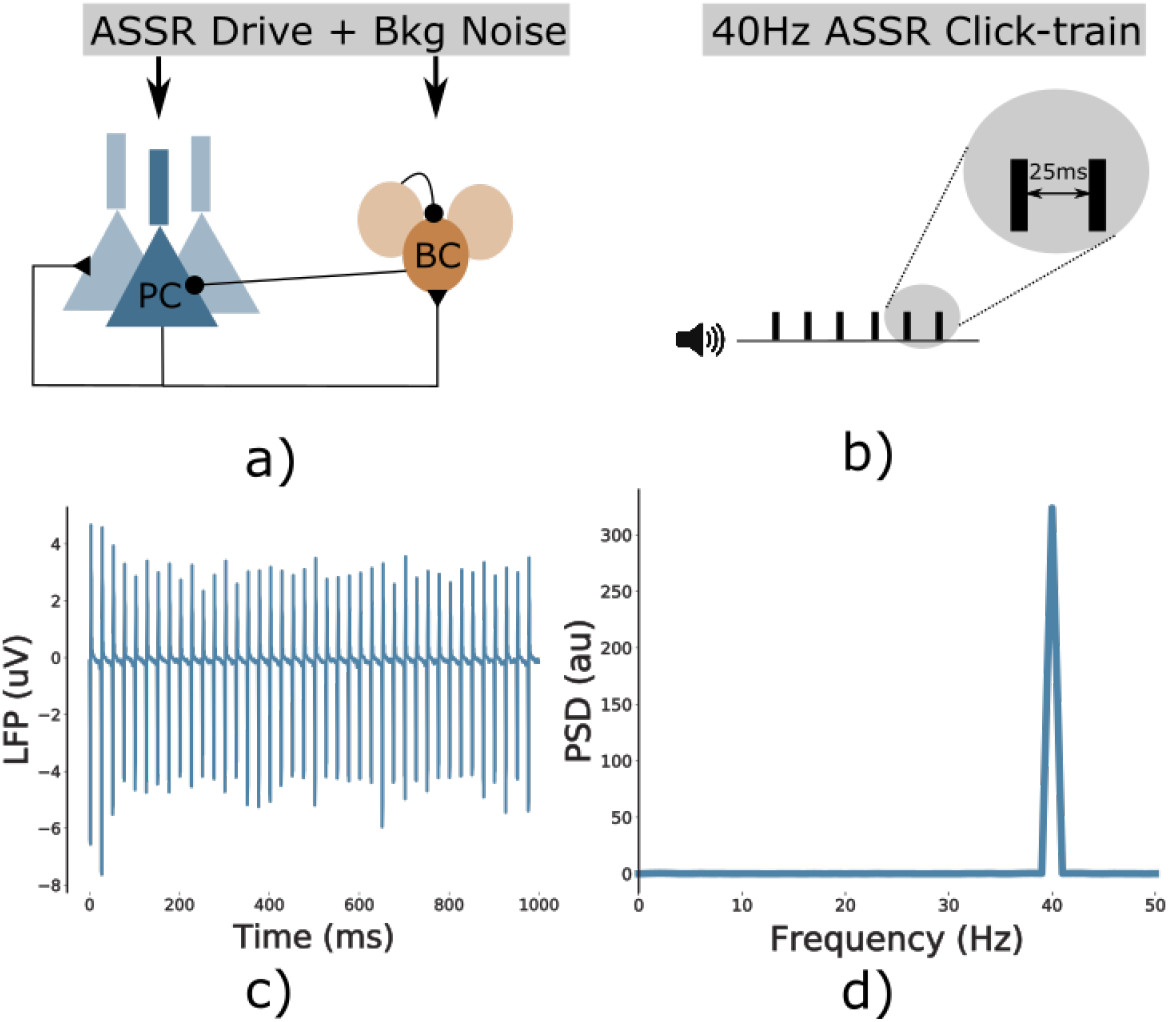
Overview. (a) Network schematic. The network consists of two interconnected populations, an excitatory population of pyramidal cells and an inhibitory population of basket cells, both receiving rhythmic ASSR drive and Poissonian background noise. (b) The 40 Hz ASSR drive is modeled as bouts of input spikes arriving at all cells simultaneously with an inter-bout interval of 25 ms, mimicking a 40 Hz ASSR click-train paradigm. (c) Example LFP signal of the control network in response to the ASSR drive, showing strong 40 Hz entrainment. (d) Power spectral density of the signal from c), again confirming that the network follows the 40 Hz click train rhythm.

**Figure 2:**
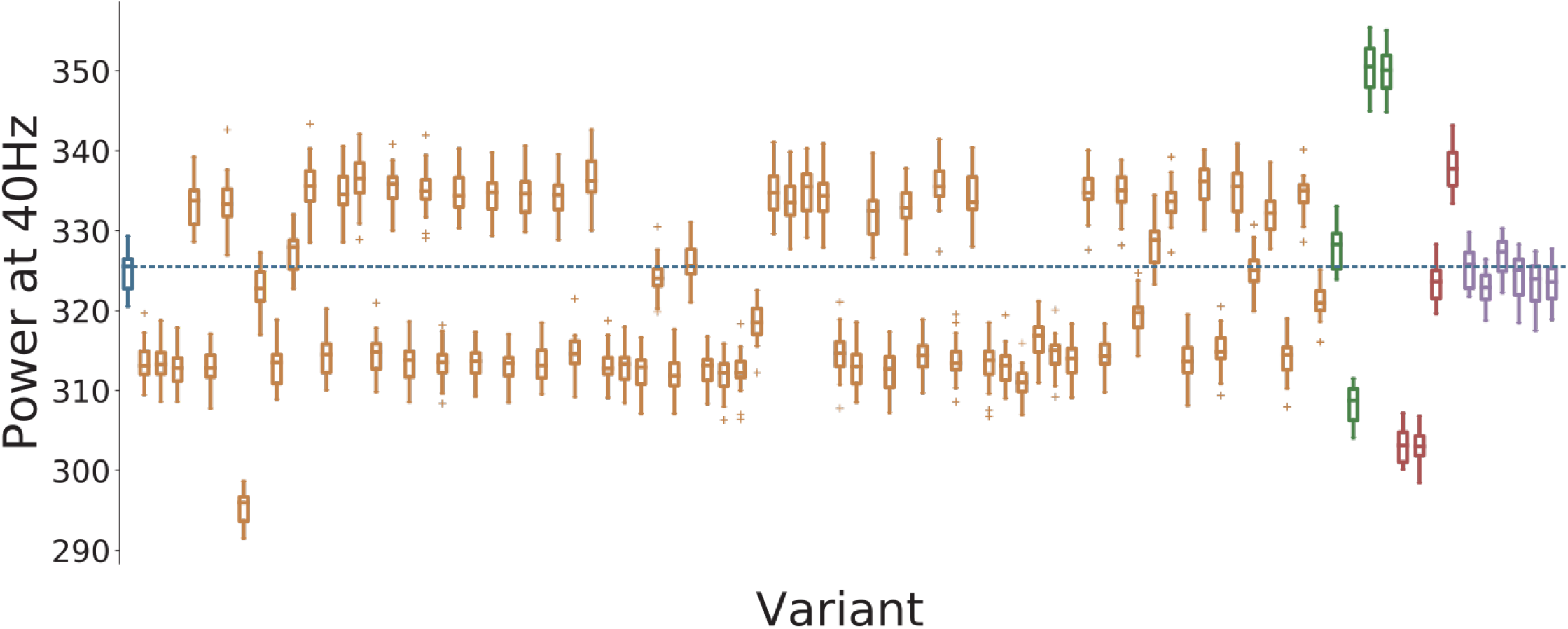
Overview of the 4040 measure for all variants. Control network in blue, *Ca*^2+^ channel variants affecting Ca_*HV A*_ in orange, *Ca*^2+^ channel variants affecting Ca_*LV A*_ in green, HCN variants in red, and SCN variants in purple. Solid lines represent the mean, box edges the 25 and 75 percentile, respectively, the whisker extend to 2 standard deviations and + depict outliers. The dashed blue line represents the mean of the control network.

**Figure 3:**
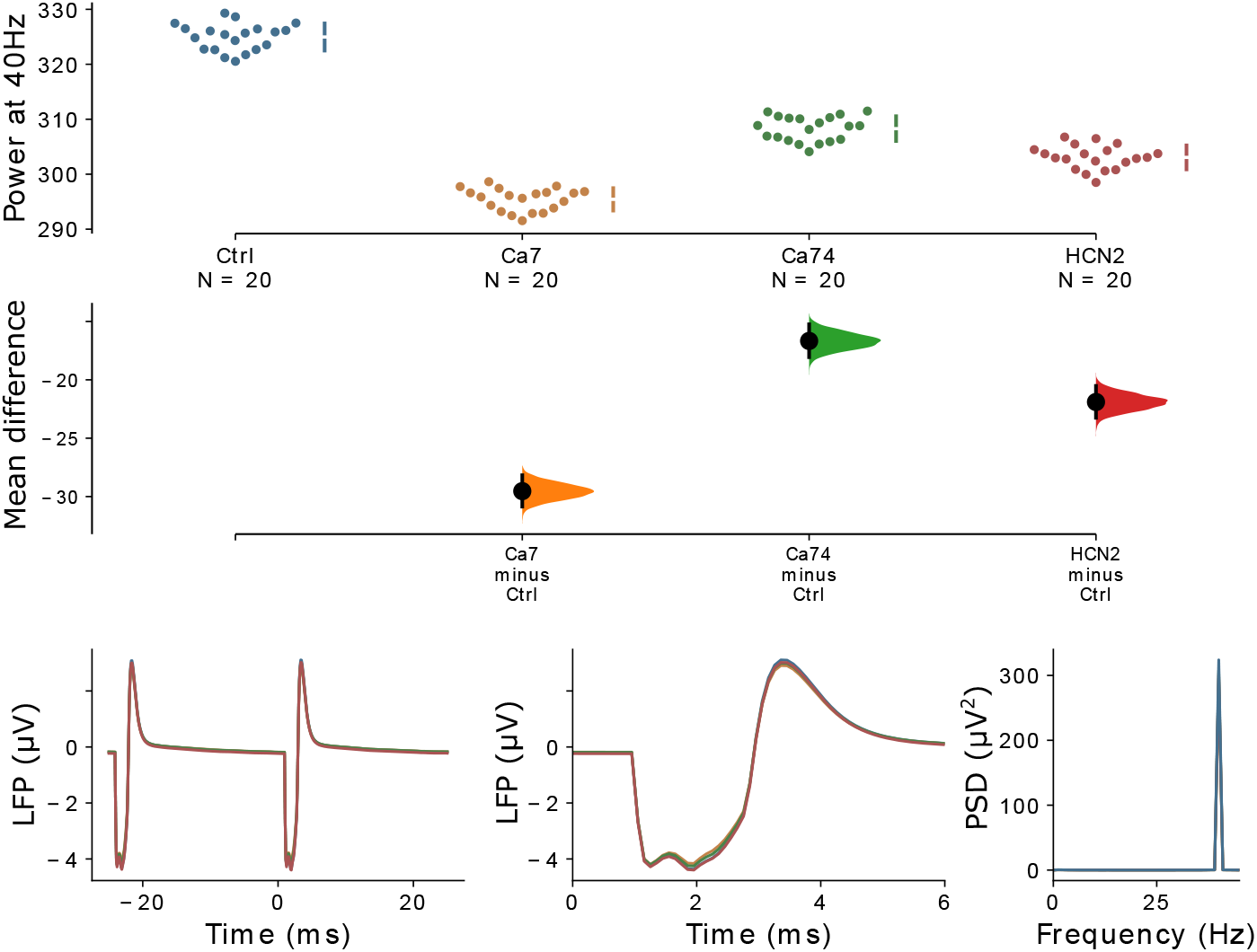
Comparison of different variants against the control. (a) The mean difference, i.e. for three comparisons (the Ca^2+^ channel model variants, Ca7 and Ca74, and the HCN model variant, HCN1-2, with the strongest gamma reduction) against the shared control are shown in the above Cumming estimation plot. The raw data is plotted on the upper axes. On the lower axes, mean differences are plotted as bootstrap sampling distributions. Each mean difference is depicted as a dot. Each 95% confidence interval is indicated by the ends of the vertical error bars. (b) The simulated LFP signal for the control network (blue), Ca7 (yellow), Ca74 (green), and HCN1-2 (red) is shown, averaged over two consecutive gamma cycles. (c) The two signals from (b) are presented with a zoom into the narrow time frame after the stimulus arrives at 0 ms. (d) The power spectral density (PSD) for the LFP signals from (b). Note that in (b)-(d) the LFP signal is first averaged over all ‘subjects’ and ‘trials’.

### 3.2 Combinations of Variants Can Produce Substantial ASSR Power Deficits

Next, we tested three different combinations of model variants (details in Supplementary Tables 2 and 3). The combinations were: (1) the combination of the two model variants with the strongest gamma reduction from the single model variant trials (Ca7 affecting gene CACNA1C and HCN1-2 affecting HCN1; this combination will be referred to as *Comb1*); (2) the model variants from (1) but additionally one more model variant with a moderate gamma reduction (Ca74 affecting CACNA1I gene, this combination will be referred to as *Comb2*); (3) the model variants from Comb2 plus an additional model variant with a moderate gamma reduction (HCN1-1 affecting the HCN1 gene; this combination will be referred to as *Comb3*)). In the last case, when combining model variants of the same gene, we assume a linear superposition of the effects of the single model variants on the parameters of the ion channels. Note that this is a simplistic assumption and that there could potentially be nonlinear interactions between different variants of the same gene. However, actual experimental data on this relationship are currently not available. We found that combining model variants further increased their effect on evoked gamma power and a combination of only a few variants already had a strong impact on gamma power (Figure 4 and Supplementary Table S4). Overall, the effects of combining model variants were additive, e.g. combining the two model variants Ca7 and HCN1-2 which individually have a mean difference of −29.5 and −21.9 results in a combination with a mean difference of −49.7 which is roughly the sum of the two individual mean differences (see also Supplementary Table 4).

**Figure 4:**
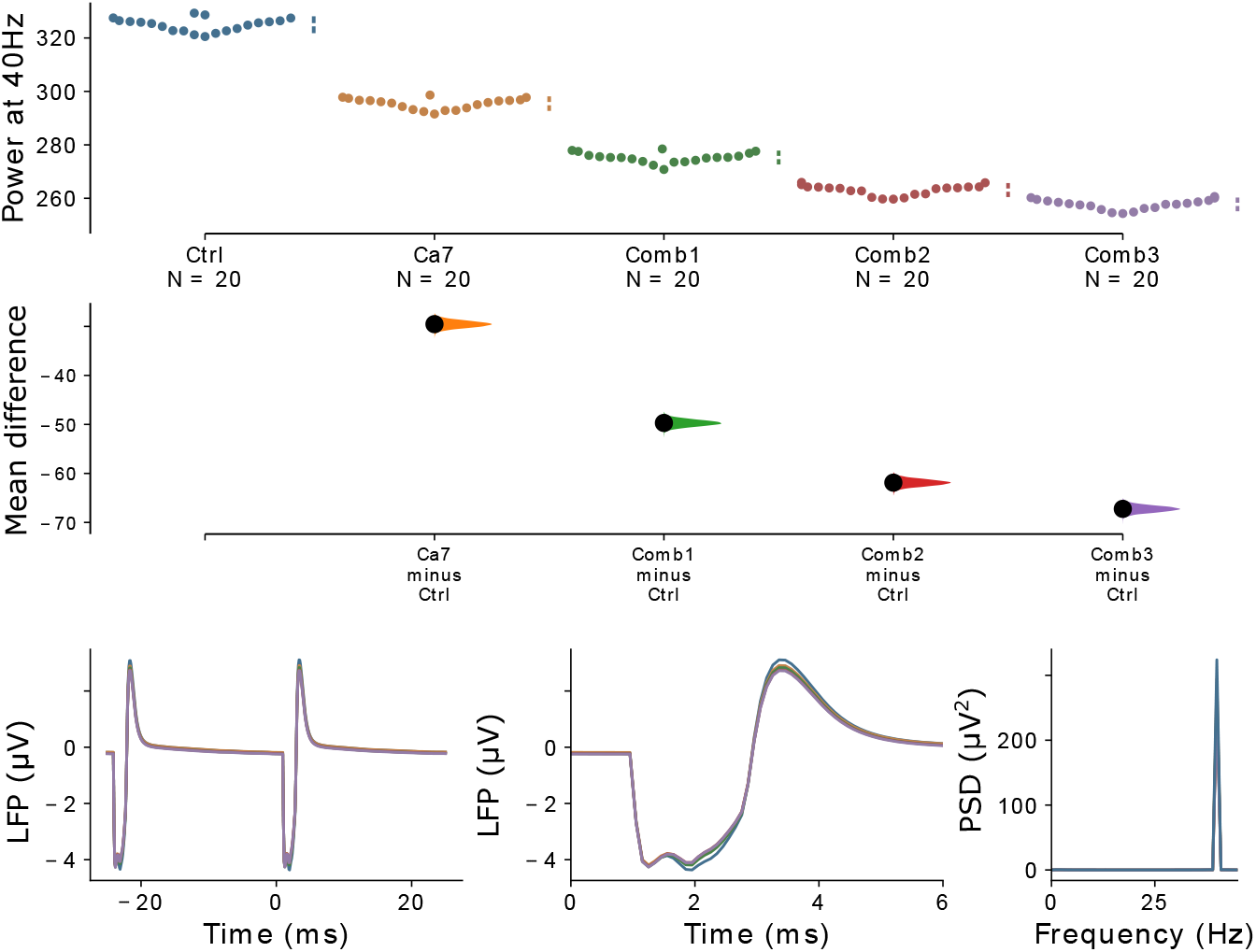
Comparison of different combinations of variants against the control. The mean difference for the comparisons of three variant combinations and the single variant with the strongest effect (Ca7) against the shared control are shown in the above Cumming estimation plot. The raw data is plotted on the upper axes. On the lower axes, mean differences are plotted as bootstrap sampling distributions. Each mean difference is depicted as a dot. Each 95% confidence interval is indicated by the ends of the vertical error bars.

### 3.3 The Effects of Variant Combinations Are Comparable to the Effects of Synaptic Alterations

Changes to the GABAergic system have been proposed to explain the reduction in evoked gamma power [32] and modelling work has demonstrated that both, a reduction in GABA levels in schizophrenia patients [89, 70] and an increase in decay times at GABAergic synapses [104, 70], can lead to reduced ASSR gamma power. On the one hand, a decrease in inhibition due to reduced GABA levels decreases the precise inhibitory control over pyramidal cell firing necessary for a strong gamma rhythm. On the other hand, an increase in GABAergic decay times, while increasing inhibition, can lead to suppression of pyramidal cell firing every second gamma cycle (a phenomenon called ‘beat-skipping’) and thus strongly reduce gamma power. Therefore, we compared the model variant effects to the effects of these two changes to the GABAergic system of the network. Specifically, we implemented a 25% reduction of the maximum conductance of GABAergic synapses (see e.g. [70]) and an increase of the decay time of inhibitory postsynaptic currents (IPSCs) at GABAergic synapses from 8 ms to 25 ms (e.g. [104, 70]). These two conditions will be referred to as *Gmax* and *IPSC*. Comparing the effects of the model variants and their combinations to these two changes allowed us to judge the relative size of their effect.

Figure 5 shows that, while even the strongest individual model variants only result in moderate reductions of gamma power, combinations of several model variants can produce strong gamma reduction comparable to a strong reduction of GABA levels (see also Supplementary Table 4). Interestingly, the increase in GABAergic decay times produced a substantially stronger reduction of gamma power, mainly due to the emergence of a beat-skipping behaviour, which shifted the power from the 40 Hz gamma band to the subharmonic 20 Hz beta band (see Supplementary Figure 8). Nevertheless, these results demonstrate that combinations of model variants, influencing ionic channels and single cell excitability, can potentially have strong effects on gamma entrainment.

**Figure 5:**
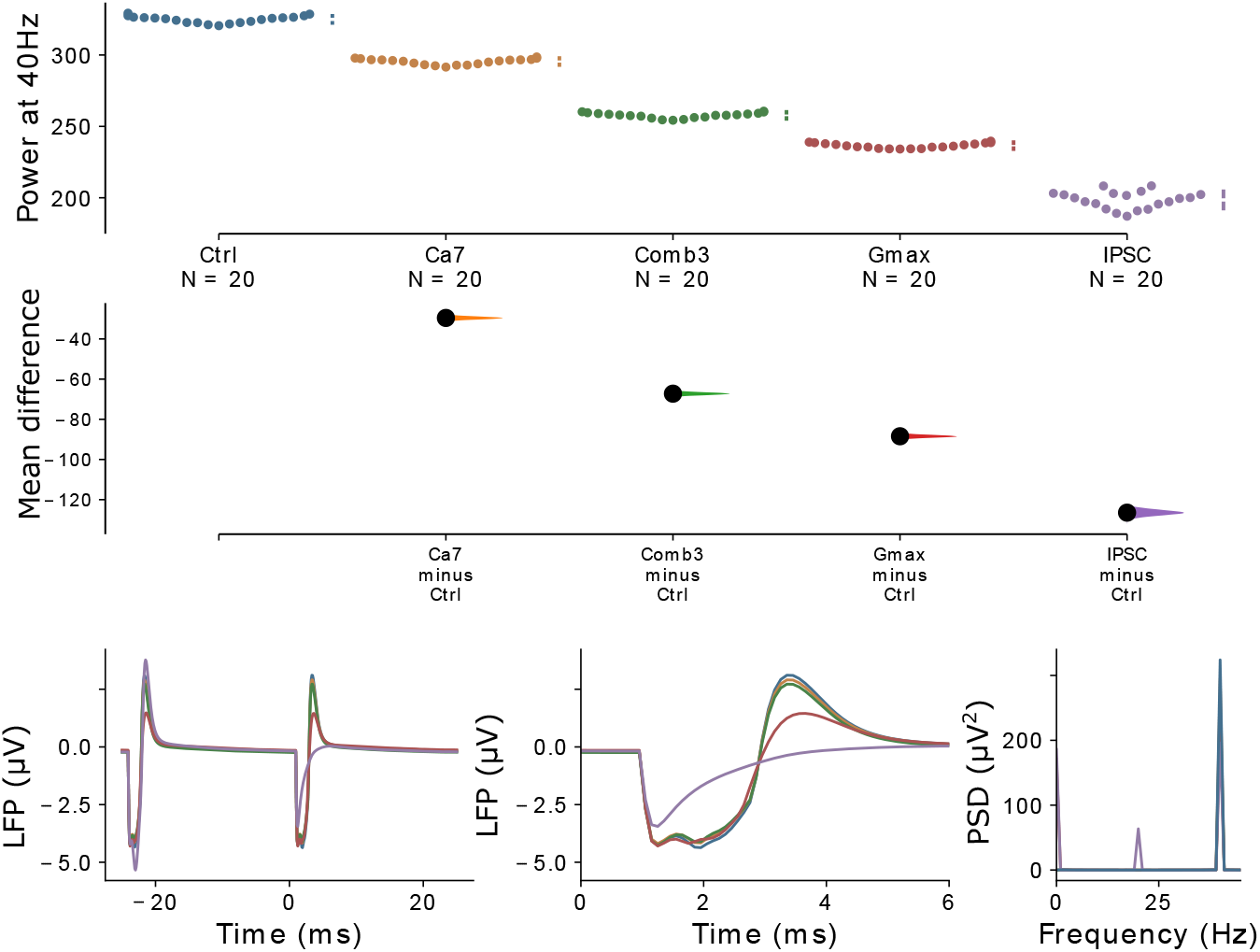
Comparison of model variants against synaptic alterations. The mean difference for the comparisons of the single model variant and the model variant combination with the strongest gamma reduction together with the two synaptic mechanisms, *Gmax* and *IPSC*, against the shared control are shown in the above Cumming estimation plot. The raw data is plotted on the upper axes. On the lower axes, mean differences are plotted as bootstrap sampling distributions. Each mean difference is depicted as a dot. Each 95% confidence interval is indicated by the ends of the vertical error bars.

### 3.4 Genetic Variants Do not Affect the Inter-Trial Phase Coherence

Besides strong reductions of evoked power, many studies report a decrease in inter-trial phase coherence (ITPC), a measure of how aligned the phase angles of the signal over individual trials are, during gamma entrainment in patients with schizophrenia (e.g. [51, 49], see Thune et al. [95] for a review). Therefore, we also calculated the ITPC for each single model variant and compared them to the two synaptic conditions from before (see 2). We found that none of the model variants altered the phase coherence substantially, while both synaptic conditions strongly decreased ITPC (Figure 6). This means that while both synaptic conditions reduce phase coherence, i.e. desynchronize the network, the reduction in gamma power resulting from the model variants seems to come solely from a reduction in amplitude of the entrained oscillations, leaving the temporal precision intact. Furthermore, this suggests that, while genetic variants of ion channel-encoding genes might contribute to the reductions in evoked gamma power found in patients with schizophrenia, it is unlikely that they play an important role in the emergence of decreases in phase coherence and the underlying desynchronization of activity in the network.

**Figure 6:**
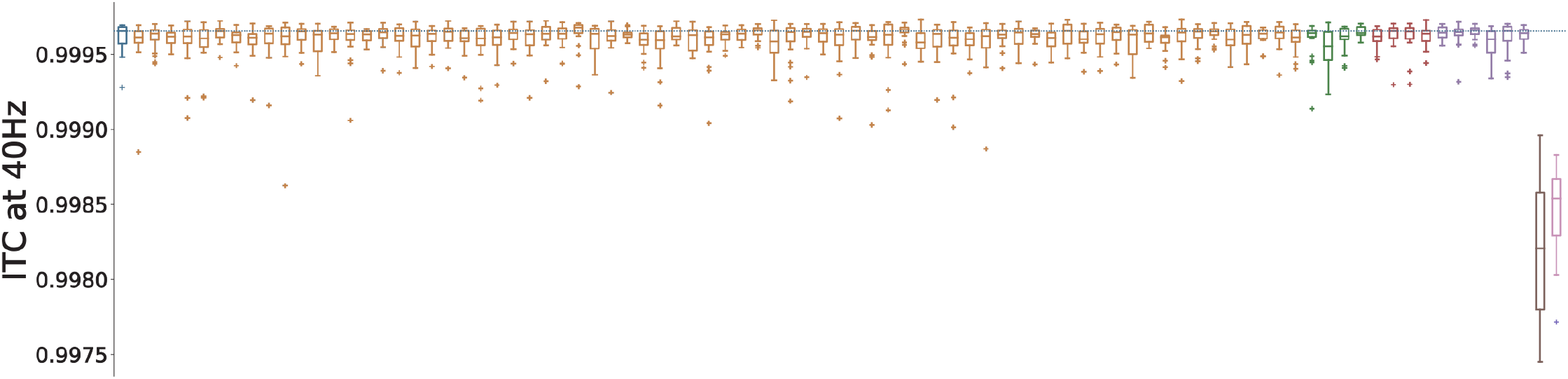
Overview of 4040 inter-trial coherence for all variants. Control network in blue, *Ca*^2+^ channel model variants affecting Ca_*HV* *A*_ in yellow, *Ca*^2+^ channel model variants affecting Ca_*LV* *A*_ in green, HCN model variants in red, SCN model variants in purple, the reduction of *g_max_* in brown and increased IPSC times in steel pink. Solid line represents the mean, box edges the 25 and 75 percentile, respectively, the whisker extend to 2 standard deviations and + depict outliers. the dashed blue line depicts the mean of the control network.

### 3.5 Correlations to Pre-Pulse Inhibition and Delta Resonance

Lastly, we compared the effect of the genetic variants with their effect on other potential biomarkers for SCZ from our earlier studies [63]. We calculated Pearson correlation coefficients between the ratio of gamma reduction and the delta resonance power and the pre-pulse inhibition (PPI) thresholds for all 86 model variants (see Supplementary Methods for details). We found a strong positive correlation between gamma power and delta resonance power (Pearson r=0.683, p<0.0001) and moderate negative correlation between gamma power and PPI threshold (Pearson r=−0.365, p<0.0001).

## 4 Discussion

In this modelling study we showed that changes to the kinetics of voltage-gated ion channels due to SCZ-associated common variants of their encoding genes led to decreases of evoked gamma power, one of the most frequently reported electrophysiological phenotypes in SCZ (Figures 2 and 3). We further demonstrated that combinations of model variants produced larger decreases in evoked gamma power (Figure 4) and that these decreases were comparable to alterations at the synaptic level (Figure 5), which are more commonly associated with changes of evoked oscillatory power in the gamma range [104, 70]. Interestingly, we found that the genetic variants, opposed to the synaptic alterations, did not change the inter-trial phase coherence, a measure of synchronization. The reductions in evoked gamma power were solely due to reductions in the amplitude of the oscillations (Figure 5).

The genetic variants that decreased evoked gamma power identified in this study mostly decreased the excitability of the layer 5 pyramidal cell, as found in earlier work [60], and, therefore, led to smaller amplitudes of the evoked LFP signals. Not surprisingly, there was not much overlap with the variants we previously found to substantially increase network delta oscillations and to reduce single cell PPI [63]. We further analysed this by correlating gamma power with delta resonance power and PPI threshold from this previous study, respectively. Here, we found a strong positive correlation between gamma and delta resonance power and a moderate negative correlation with PPI thresholds. This suggests that variants decreasing gamma power also lead to lower delta resonance power and larger PPI thresholds. While robust evidence for decreased PPI thresholds in SCZ exists [96], the findings on delta power show mixed results. Several studies find increased delta power in patients [72, 7, 41, 36, 24], however, decreased delta oscillation power has also been reported [23, 26]. Nonetheless, as also described earlier, it seems unlikely that gamma power reduction, delta power changes and PPI threshold decrease are solely caused by genetic variants affecting ionic channels. Gamma reductions most certainly are at least partially attributable to synaptic alterations as explained earlier. Nevertheless, the variants modelled here might play an important role in gamma reduction in subpopulations of patients and contribute to the large heterogeneity observed in patients with schizophrenia.

Previous models of gamma range oscillatory deficits in SCZ have mainly focused on changes at the synaptic level, such as changes of GABAergic synapses from PV^+^ interneurons onto pyramidal cells or other PV^+^ interneurons [70, 71, 104, 86, 89, 106], changes of glutamatergic excitation of PV^+^ interneurons through NMDA receptors, [46, 86] or changes of spine density at pyramidal cells [86]. As mentioned in the Results section, there are two main consequences of changes to the GABAergic system that have been the focus of previous modelling studies: 1) A reduction of the peak amplitude of the IPSC and 2) a prolongation of IPSC decay times. A reduction of the peak IPSC amplitude has been shown to significantly reduce evoked gamma power but to leave beta power intact [70, 86, 89]. On the other hand, an increase of IPSC decay time, while also substantially reducing evoked gamma power, has been shown to increase power in the beta band [104, 70] and most probably exerts its effect through PV^+^ basket cells [71]. Kirli et al. [46, 48] demonstrated that lower gamma band power was present at both low and high NMDA conductance levels with optimal synchronization occurring at intermediate conductance levels. In another study, Siekmeier and van Maanen [86] showed that modest reductions in NMDA system function and dendritic spine density led to a robust reduction of gamma power. However, they also found that greater NMDA hypofunction along with low level GABA system dysregulation substantially decreased gamma power, highlighting the multifactoriality of underlying alterations. In addition, dopaminergic modulation of local circuits has also been shown to affect gamma synchronization through modulation of K^+^ currents and NMDA conductances [47]. Overall we must note that, as demonstrated for models of gamma ASSRs in particular [86, 70] but also other models of psychiatric disorders [78] and models of healthy local circuits [81] in general, many different parameter combinations might produce similar network level behaviour. Therefore, it is very important to explore the interaction of alterations of different systems. Moreover, to constrain the models further, these interactions should be tested against different paradigms, such as spontaneous and evoked oscillations. The computational model presented here offers an ideal starting point for such an effort, since it allows for the integration of the most crucial factors contributing to gamma band oscillatory deficits in schizophrenia. Beyond incorporating variants of ion channel encoding genes and alterations of GABAergic synapses, extensions of the model could include the integration of NMDAR hypofunction via its NMDARs and the integration of changes to dopaminergic neuromodulation via its K^+^ channels, for example as in [47].

A crucial part of the overall approach used in this study is the dampening of the effect of literature-derived model variants by a downscaling of the parameter changes. Typically, the literature-derived model variants very strongly changed the physiology of the studied cell (see also [60]) and, therefore, were not representative of common SNP-like variants. Overall, single SNP-like variants, which are known to be numerous and to occur frequently in the healthy population [83], are assumed to have small phenotypic effects, either on the single cell or on the systems level. Nevertheless, we note that rare variants with large effects associated with schizophrenia also exist (see e.g. [2]). Additionally, we have shown in previous work on increased delta oscillations due to genetic variants that model variants derived from gene expression data largely result in similar changes to the network model [63].

GWAS studies, such as Ripke et al. [83], have identified numerous variants associated with psychiatric disorders, however, we know very little about their functional effects. As we have argued before [63, 62], the modelling framework presented in this study is ideally suited to build hypotheses about their effects and to make experimentally testable predictions. To be more specific, the biophysically detailed model used here can provide very specific associations between genetic variants and phenotypes, while explicitly revealing the cellular properties through which the two are mechanistically linked. This goes well beyond the purely statistical associations that standard genetics approaches produce.

### Limitations and future directions

The analysis presented here is based on the model of a thick-tufted layer 5 pyramidal cell, which accurately reproduces many active and passive electrophysiological features of these cells [37], and the effects of the genetic variants were implemented as changes to the kinetics of the underlying ion channels of the model. Therefore, our approach here rests on the assumption that the model faithfully reproduces the ion channel dynamics of layer 5 pyramidal cells. While the fitting of the model did not include the replication of activity in the presence of ionchannel blockers [58], the model’s ion channel composition is largely consistent with that of other models. Almog et al. present a model of a layer 5 pyramidal cell where the set of channels is partly overlapping [1]. Although the contributions of the ion channels to model behaviour differ slightly, with the persistent K^+^ having a larger role, the two models mostly conform with each other [1, 64]. Papoutsi et al. [76] also present a model of layer 5 pyramidal cells showing similar interactions between voltage-gated Ca^2+^ channels and the Ca^2+^-dependent afterhyperpolarization (AHP) current as in the model of Hay et al. we used here and further underpin the validity of our model assumptions.

While in this study we have focused on the effect of genetic variants on ion channel dynamics, future studies should also explore their role in schizophrenia-associated changes to synaptic receptors, especially GABA and NMDA receptors. These are, as mentioned before, more traditionally associated with gamma band deficits in the disorder [32, 33]. Furthermore, the approach outlined in this study could also be extended to study the effect of genetic variants on intracellular signaling cascades involved in plasticity [22, 61]. A major challenge for the field, however, will be to incorporate immune pathways, which have been strongly indicated by recent GWASs [83, 101], into models of schizophrenia pathophysiology.

Nevertheless, the model developed in this study can already be used to complement more traditional approaches to identify potential treatment targets. Current medications with known courses of action can be included into the model and their effects on gamma band oscillations can subsequently be assessed (see [86] for an excellent implementation of such an approach). Furthermore, a search for novel therapeutic targets can be conducted through a more exploratory analysis of the effect of parameters on the model behaviour (also see [86]). Additionally, such an approach is not limited to pharmacological interventions since transcranial electric or magnetic stimulation can easily be incorporated (as demonstrated in other modeling studies such as [10, 85, 82]); this is, of course, not restricted to schizophrenia but can be applied to psychiatric disorders in general.

In conclusion, our work represents a step towards the integration of the wealth of genetic data on psychiatric disorders into biophysically detailed models of biomarkers with great potential to unravel underlying polygenic cellular-based mechanisms. Furthermore, the approach offers an ideal test ground for the identification of novel therapeutic strategies, such as pharmaceutical interventions or electrical stimulation.

## Supplementary Material

### Supplementary Methods

#### Ion channels and their genetic etiology

The single cell model used to model the pyramidal cells in our network is based on a detailed, multi-compartment Hodgkin-Huxley type model of layer 5 pyramidal cells with a reconstructed morphology [37]. However, the very high morphological detail (196 compartments), and thus very high computational complexity, of this neuron model renders it unsuitable for the network model analysis performed in this study. Therefore, we employed a reduced version of this model, where passive parameters and ion channel and Ca^2+^ dynamics were fitted to the original model using a multi-step fitting procedure [59]. The reduced model, analogous to the original model, contains the following ionic currents: Fast inactivating Na^+^ current (*I*_Nat_), persistent Na^+^ current (*I*_Nap_), non-specific cation current (*I*_h_), muscarinic K^+^ current (*I*_m_), slow inactivating K^+^ current (*I*_Kp_), fast inactivating K^+^ current (*I*_Kt_), fast non-inactivating K^+^ current (*I*_Kv3.1_), high-voltage-activated Ca^2+^ current (*I*_CaHVA_), low-voltage-activated Ca^2+^ current (*I*_CaLVA_), small-conductance Ca^2+^-activated K^+^ current (*I*_SK_), and the passive leak current (*I*_leak_).

Exact information on which ion channel subunits contribute to the above mentioned currents in layer 5 pyramidal cells is still missing. However, the expression of different ion channel subunits in these cells has been studied extensively, and indications can be drawn from these studies. Christophe et al. [16] found the mRNA of the ion channel-encoding genes *KCNA2*, *KCND2*, *KCND3*, *CACNA1A*, *CACNA1B*, *CACNA1C*, *CACNA1D*, *CACNA1E*, *CACNA1G*, *CACNA1H*, *CACNA1I*, *HCN1*, and *HCN2* expressed in postnatal rat neocortices at different stages of development. Of these genes, it is known that*CACNA1A*, *CACNA1B*, *CACNA1C*, and *CACNA1D* contribute to *I*_CaHVA_, *CACNA1G*, *CACNA1H*, and *CACNA1I* to *I*_CaLVA_, while the genes *KCNA2*, *KCND2*, and *KCND3* might contribute to the slow *I*_Kp_ current.

In a different study, the genes *SCN1A*, *SCN2A*, *SCN3A*, and *SCN6A* were found to be expressed in layer 5 pyramidal cells [110]. While several of the genes encoding the *α* subunits (SCN1A, SCN2A and SCN3A) are tetrodotoxin-sensitive [12] and hence form both the transient (*I*_*Nat*_) and persistent (*I_Nap_*) Na^+^ currents, their contribution to *I*_*Nat*_ or *I_Nap_* might be dependent on modulatory subunits [56].

#### Description of the Single Cell Models

##### Layer 5 pyramidal cells

The layer 5 pyramidal cell model is a multi-compartment Hodgkin-Huxley type model and the membrane potential dynamics can be described by

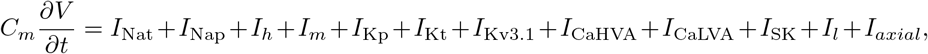

where the different currents can be written as the product of an activation and an inactivation variable

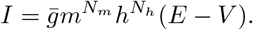

Here, *m* and *h* are the activation and inactivation variables, *N_m_* and *N_h_* their sensitivities, 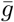 the maximal ionic conductance and *E* the ionic reversal potential. Na+ and K+ have a fixed reversal potential of *E*_*Na*_ = 50 mV and *E*_*Na*_ = −85 mV, respectively, while the reversal potential of Ca^2+^ depends on the intracellular [Ca^2+^]. Furthermore, the dynamics of activation and inactivation are defined by

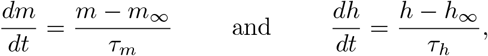

where *m*_∞_, *h*_∞_, *τ_m_*, and *τ_h_* are functions of the membrane potential *V*. In our case, *m*_∞_ and *h*_∞_ usually have a sigmoidal shape, where the half-activation and half-inactivation voltages are determined by one or more parameters each (depending on the ion channel). These parameters are written as *V*_offm∗_ and *V*_offh∗_, where stands for further specifications in case there are multiple parameters affecting them. Analogously, the parameters *V*_slom∗_ and *V*_sloh∗_ affect the slopes of the (in)activation curves, and parameters *τ*_m∗_ and *τ*_offh∗_ the time constants. All ionic currents, except for the activation of *I*_SK_, are described in this way. The activation of *I*_SK_, on the other hand, only depends on the intracellular [Ca^2+^], through a sigmoidal function with a half-activation concentration parameter *c*_off_ and a slope parameter *c*_slo_. Lastly, the dynamics of the intracellular [Ca^2+^] is described by

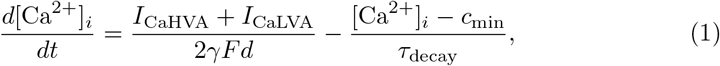

where *I*_CaHVA_ and *I*_CaLVA_ are the high and low-voltage activated Ca^2+^ currents entering the considered cell segment, *γ* represents the fraction of Ca^2+^ ions entering the cell that contribute to the intracellular [Ca^2+^], *F* the Faraday constant, *d* is the depth of the sub-membrane layer considered for calculation of concentration, *c*_min_ the resting intracellular [Ca^2+^], and *τ*_decay_ is the decay time constant of the intracellular [Ca^2+^].

As mentioned earlier, in this study we employed the reduced model presented in [59], consisting of four compartments: the soma, the apical trunk, the apical tuft, and the basal dendrite. the maximal conductance values 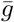 were different for different compartments and are subindexed as follows: *s* refers to the soma, *a*_0_ to the apical trunk, *a*_1_ to the apical tuft, and *b* to the basal dendrite. All values not shown are set to 0.

**Fast inactivating Na**^+^ **current, *I***_Nat_

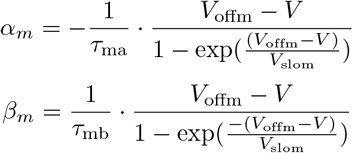

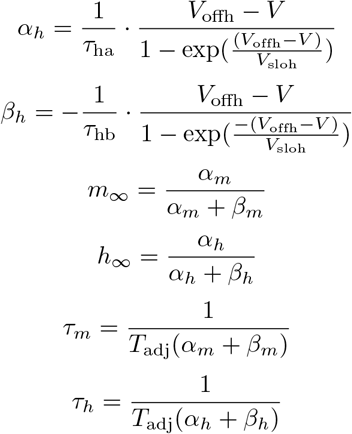

*V*_offm_ = −38 mV, *V*_offh_ = −66 mV, *V*_slom_ = 6.0 mV, *V*_sloh_ = 6.0 mV, *τ*_ma_ = 5.49 ms, *τ*_mb_ = 8.06 ms, *τ*_ha_ = 66.67 ms, *τ*_hb_ = 66.67 ms, 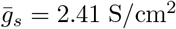,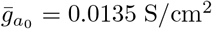, 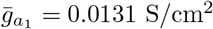 *N*_*m*_ = 3, *N*_*h*_ = 1

**Persistent Na**^+^ **current**, *I*_Nap_

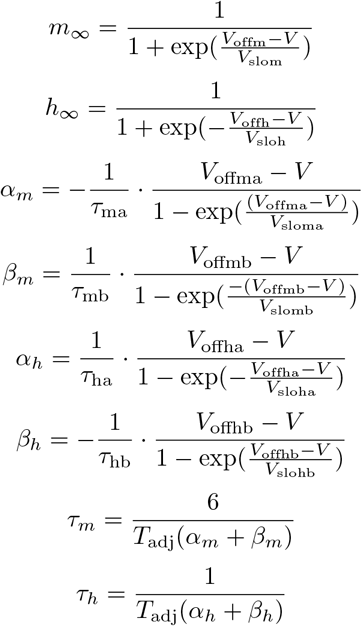

*V*_offm_ = −52.6 mV, *V*_slom_ = 4.6 mV, *V*_offma_ = −38 mV, *V*_offmb_ = −38 mV, *V*_sloma_ = 6.0 mV, *V*_slomb_ = 6.0 mV, *τ*_ma_ = 5.49 ms, *τ*_mb_ = 8.06 ms, *V*_offh_ = −48.8 mV, *V*_sloh_ = 10.0 mV, *V*_offha_ = −17 mV, *V*_offhb_ = −64.4 mV, *V*_sloha_ = 4.63 mV, *V*_slohb_ = 2.63 mV, *τ*_ha_ = 347222.2 ms, *τ*_ha_ = 144092.2 ms, 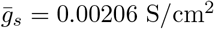, *N_m_* = 3, *N_h_* = 1

**Non-specific cation current, *I***_h_

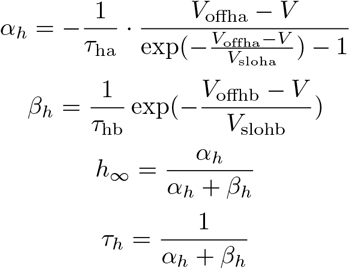

*E* = −45.0 mV, *V*_offha_ = −154.9 mV, *V*_sloha_ = 11.9 mV, *τ*_ha_ = 155.52 ms, *V*_offhb_ = 0.0 mV, *V*_slohb_ = 33.1 mV, *τ*_hb_ = 5.18 ms, 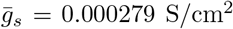,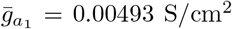, 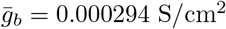, *N_m_* = 0, *N*_1_ = 0

**Muscarinic K**^+^ **current, *I***_m_

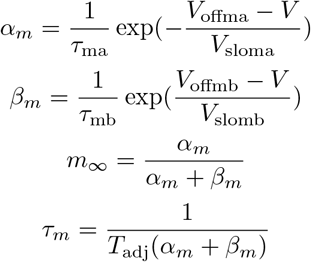

*V*_offma_ = −35 mV, *V*_sloma_ = 10 mV, *τ*_ma_ = 303.03 ms, *V*_offmb_ = −35 mV, *V*_slomb_ = 10 mV, *τ*_mb_ = 303.03 ms, 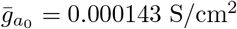, 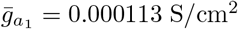, *N_m_* = 1, *N_h_* = 0

**Slow inactivating K**^+^ **current, *I***_Kp_

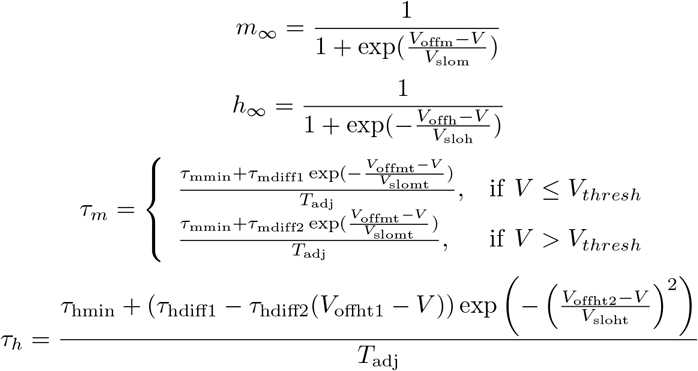

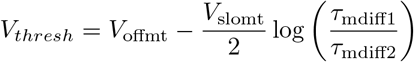

*V*_offm_ = −11 mV, *V*_slom_ = 12 mV, *V*_offmt_ = −10 mV, *V*_slomt_ = 38.46 mV, *τ*_mmin_ = 1.25 ms, *τ*_mdiff1_ = 175.03 ms, *τ*_mdiff2_ = 13 ms, *V*_offh_ = −64 mV, *V*_sloh_ = 11 mV, *V*_offht1_ = −65 mV, *V*_offht2_ = − 85 mV, *V*_sloht_ = 48 mV, *τ*_hmin_ = 360 ms, *τ*_hdiff1_ = 1010 ms, *τ*_hdiff2_ = 24 ms/mV, 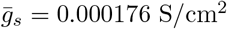, *N_m_* = 2, *N_h_* = 1

**Fast inactivating K**^+^ **current, *I***_Kt_

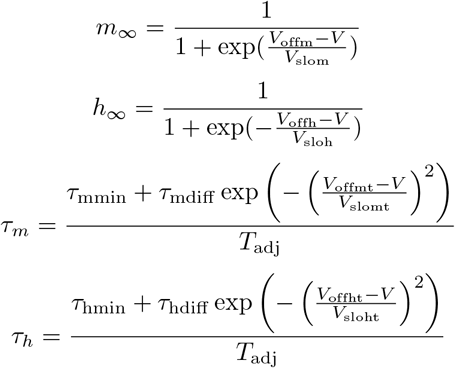

*V*_offm_ = −10 mV, *V*_slom_ = 19 mV, *V*_offh_ = −76 mV, *V*_sloh_ = 10 mV, *V*_offmt_ = −81 mV, *V*_slomt_ = 59 mV, *τ*_mmin_ = 0.34 ms, *τ*_mdiff_ = 0.92 ms, *V*_offht_ = − 83 mV, *V*_sloht_ = 23 mV, *τ*_hmin_ = 8 ms, *τ*_hdiff_ = 49 ms, 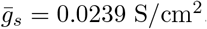, *N_m_* = 4, *N_h_* = 1

**Fast, non inactivating K**^+^ **current, *I***_Kv3.1_

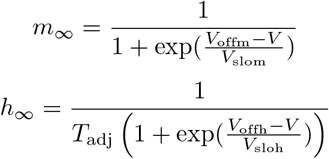

*V*_offma_ = 18.7 mV, *V*_offmt_ = −46.56 mV, *V*_sloma_ = 9.7 mV, *V*_slomt_ = 44.14 mV, *τ*_mmax_ = 4.0 ms, 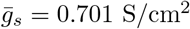, 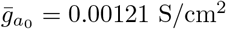, *N_m_* = 1, *N_h_* = 0

**High-voltage-activated Ca**^2+^ **current, *I***_CaHVA_

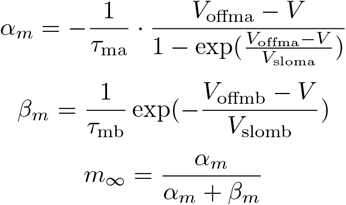

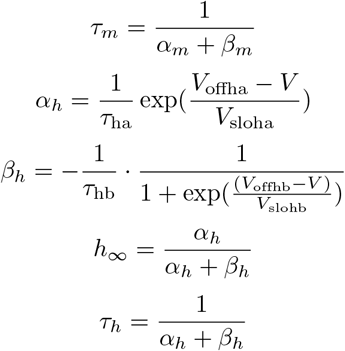

*V*_offma_ = −27 mV, *V*_offmb_ = −75 mV, *V*_offha_ = −13 mV, *V*_offhb_ = −15 mV, *V*_sloma_ = 3.8 mV, *V*_slomb_ = 17 mV, *V*_sloha_ = 50 mV, *V*_slohb_ = 28 mV, *τ*_ma_ = 18.18 ms, *τ*_mb_ = 1.06 ms, *τ*_ha_ = 2188.18 ms, *τ*_hb_ = 153.85 ms, 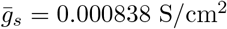, 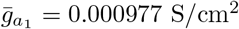, *N_m_* = 2, *N_h_* = 1

**Low-voltage-activated Ca**^2+^ **current, *I***_CaLVA_

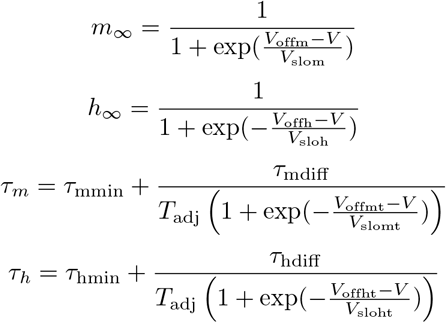

*V*_offma_ = −40.0 mV, *V*_offmt_ = −35.0 mV, *V*_offha_ = −90.0 mV, *V*_offht_ = −50.0 mV, *V*_sloma_ = 6.0 mV, *V*_slomt_ = 5.0 mV, *V*_sloha_ = 6.4 mV, *V*_sloht_ = 7.0 mV, *τ*_mmin_ = 5.0 ms, *τ*_mdiff_ = 20.0 ms, *τ*_hmin_ = 20.0 ms, *τ*_hdiff_ = 50.0 ms, 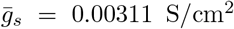, 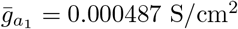, *N_m_* = 2, *N_h_* = 1

**Small-conductance Ca**^2+^**-activated K**^+^ **current, *I***_SK_

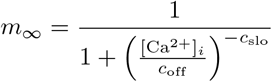

*c*_off_ = 0.00043 mM, *c*_slo_ = 4.8, 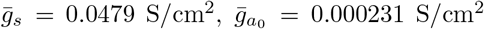, 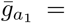, 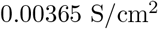, *N_m_* = 1, *N_h_* = 1

**Leak current, *I***_leak_

*E* = −90 mV, 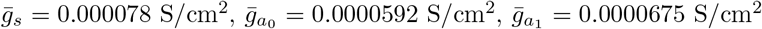, 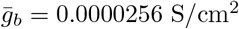, *N_m_* = 0, *N_h_* = 0

##### Intracellular [Ca^2+^] dynamics

The intracellular Ca^2+^ concentration follows Equation 1 with the following model parameters: *γ*_*s*_ = 0.0005, 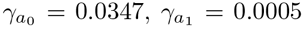, *τ*_decay,*s*_ = 488 ms, 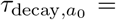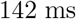, 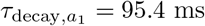, *d* = 0.1 μm, *c*_min_ = 10^−4^ mM

**Temperature adjustment factor**: 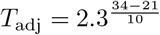

##### Inhibitory interneurons

Fast-spiking interneurons were also modelled as multi-compartment Hodgkin-Huxley type neurons. The model was taken from Vierling-Claassen et al. [103] which can be found in ModelDB (https://senselab.med.yale.edu/modeldb/ShowModel?model=141273). The evolution of its membrane potential over time is governed by the following differential equation

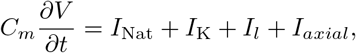

where the currents are modelled with the same formalisms as for the pyramidal cells. The two ionic currents *I_Nat_* and *I_K_* were modelled as in [57].The basket cell model consisted of 16 compartments: the soma, and 15 dendritic compartments. The maximal conductance values 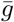 were different for different compartments and are subindexed as follows: *s* refers to the soma, *d* to the dendrite (for details see [103]). All values not shown are set to 0.

**Fast inactivating Na**^+^ **current, *I***_Nat_

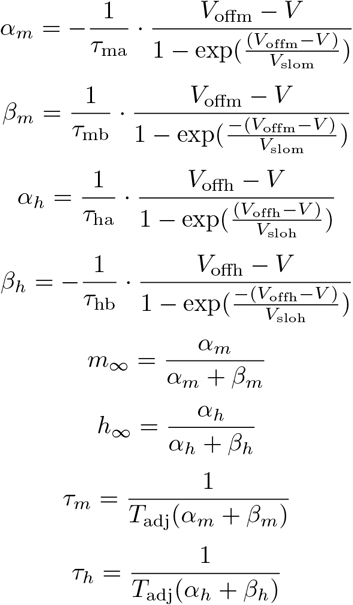

*V*_offm_ = 25 mV, *V*_offh_ = 25 mV, *V*_slom_ = 9.0 mV, *V*_sloh_ = 9.0 mV, *τ*_ma_ = 5.49 ms, *τ*_mb_ = 8.06 ms, *τ*_ha_ = 41.67 ms, *τ*_hb_ = 109.89 ms, 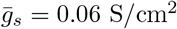,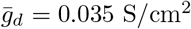, *N_m_* = 3, *N_h_* = 1

**K**^+^ **current, *I***_K_

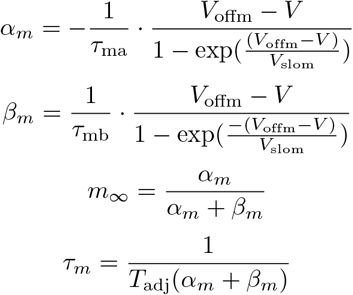

*V*_offm_ = 25 mV, *V*_offh_ = 25 mV, *τ*_ma_ = 50.0 ms, *τ*_mb_ = 500.0 ms, 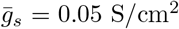, 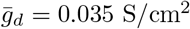, *N_m_* = 1.

**Leak current,** *I*_leak_

*E* = 73 mV, 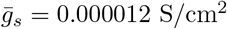, *N_m_* = 0, *N_h_* = 0

For the interneurons, Na^+^ and K^+^ have a fixed reversal potential of *E*_*Na*_ = 50 mV and *E*_*Na*_ = −85 mV.

#### Description of the Network Model

The network model consisted of 256 layer 5 pyramidal cells and 64 fast-spiking inhibitory interneurons and was implemented in NEURON [38] using the NetPyNE interface [25]. Pyramidal cells were connected to each other randomly with a probability of 0.06 (using AMPA and NMDA synaptic receptors). AMPA synaptic receptors were modelled as a double exponential function with a rise time of 0.1 ms and decay time of 3.0 ms. NMDA receptors were modelled as a first-order model including different binding and unbinding rates together with a magnesium block [20, 21]. AMPA receptors had a maximal conductance of 0.0012 nS and the NMDA receptors of 0.0006 nS, reflecting the higher contribution of AMPA receptors (see [75, 89]). Pyramidal cells were connected to inhibitory interneurons with a probability of 0.43, again using AMPA and NMDA receptors. However, for this connection type the maximal conductances for AMPA receptors, 0.0012 nS, was substantially higher than for NMDA, 0.00013, reflecting the minor role NMDAergic activation plays in the recruitment of PV^+^ interneurons [73, 89, 31]. Inhibitory interneurons formed connections with pyramidal cells and with themselves with a probability of 0.44 and 0.51, respectively. Inhibitory connections were realised using GABA_*A*_ receptors, which were also modelled as double exponential functions with a rise time of 0.5 ms and a decay time of 8.0 ms (except for the condition of prolonged inhibitory decay times, where the decay time is increased to 25.0 ms). Maximal conductance values for inhibitory connections to pyramidal cells was 0.035 nS and to themselves 0.023 nS.

The network received two types of input: 1) Poissonian noise reflecting random cortical and subcortical background activity and 2) periodic input representing auditory steady-state stimuli at gamma frequency (operationalized at 40 Hz in this study). For the random background noise we utilized NEURON’s built-in *NetStim* function with *rate* = 200, *noise* = 1.0 and *start* = 500. The parameters result in a noise stimulus reflecting a background rate of 200 Hz, that is completely random (i.e. Poissonian) which starts after 500 ms and lasts until the end of the simulation. The background *NetStim* element was connected to the pyramidal cells with a weight of 0.0325 and to the interneurons with a weight of 0.002. This setting resulted in asynchronous irregular firing of both cell populations at rates of 7.81 ± 0.07 Hz for the pyramidal cells and 6.18 ± 0.26 Hz for the inhibitory neurons, which is close to the reported spontaneous activity of ~4.9 Hz reported for auditory cortex [42]. The periodic input resembling auditory click-train stimuli was also realised with NEURON’s built-in *NetStim* function with *rate* = 40.0, *noise* = 0.0 and *start* = 1000. These parameters result in a fully periodic stimulus with an inter-event interval of 25 ms, starting after 1000 ms and ending with the end of the simulation. In summary, the network was allowed to settle for the first 500 ms, then background input was switched on and after another 500 ms stimulus input was switched on. We simulated a total of 2000 ms and used only the last 1000 ms for the analysis of steady-state entrainment.

In our network model both, the background noise and the specific connections between neurons, are based on stochastic processes. To ensure that our findings do not depend on the specific instantiations of these stochastic processes, we always performed multiple simulations with different seeds for the random number generators underlying the processes. Specifically, we simulated 20 *subjects* by choosing 20 different seeds for the random number generator underlying the formation of synaptic connections, i.e. each *subject* had the same connectivity throughout all simulations. For each *subject* we then performed 10 *trials* by choosing 10 different random seeds for the random number generator underlying the background noise. In total, we performed 200 simulations for each network configuration, with the different network configurations being the control network, 2) a single model variant, 3) a combination of different model variants, and 4) a synaptic alteration (either a reduced *g*_*max*_ or a increased *τ_decay_* at GABAergic synapses).

#### Modelling SNP-like Genetic Variants

The single cell models include Hodgkin-Huxley type description for channel activation and inactivation, and hence, changes related to certain ion-channel-encoding gene variants that have been observed in experiments can be directly attributed to a change of one or more parameters of these models. However, we only modelled changes to the pyramidal cells, which have been found to be strongly implicated by many schizophrenia-associated gene sets [88].

Here, we restricted ourselves to the following set of ion channel-encoding genes: CACNA1C, CACNA1D, CACNB2, CACNA1I, SCN1A and, HCN1. For details on the selected genes and variants see also [60, 63]. Supplementary table S1 gives all details of the different variants of these genes and in Supplementary tables S2 and S3 all details on how they were integrated into the pyramidal cell model are shown.

##### Extraction of Functional Genomics Studies

The modelling of SNP-like genetic variants is based on an extensive search of the literature on how the genes CACNA1C, CACNA1D, CACNB2, CACNA1I, SCN1A, and HCN1 change the dynamics of the underlying ion channel. We included studies performed in different animal species and different cell types because of a current lack of data for a single animal and tissue type. Our inclusion criteria were as follows:

- The study applied a genetic variant of one of the genes of interest.
- The features of the variant-expressing cell were investigated using electrophysiology or Ca^2+^ imaging.
- The change from the control cell behaviour to the variant cell behaviour could be meaningfully translated to the layer 5 pyramidal cell model.
- The observed effect of the gene variant was not purely an effect of ion channel density or expression level.

We applied the last criterion mainly because there are numerous ways that might influence such an effect [84], while the changes of ion channel dynamics are supposedly more directly dependent on the genetic encoding. Supplementary table S1 lists all the pertinent data from the included studies [50, 19, 40, 91, 94, 93, 5, 112, 79, 80, 3, 54, 17, 66, 55, 43, 14, 102, 105, 13, 65, 44, 53, 108].

In studies that reported the effects of several variant types, the ranges of possible effects are considered. In case variants considered in such studies yielded positive and negative effects, both were included, however, if the reported endpoints of the ranges were too close to the control value (i.e. less than 1 mV or less than 10% of the distance between control value and the other endpoint), only the larger deviation was considered.

##### Scaling the Gene Variants

Due to the polygenic nature of psychiatric disorders such as SCZ, it seems reasonable to assume that the disorder is not caused by a single SCZ-associated SNP but rather by a combination of sufficiently many (cf. [52]). Therefore, we followed earlier approaches [60, 63] to scale down the effect of single variants. We proceeded as follows. If a variant of the model (as shown in Supplementary Table changed the response of the neuron too strongly, we scaled down the effect of the genetic variant until the differences between control and variant model neuron were within a given bound. Specifically, down-scaling was performed until the following five criteria were fulfilled ([60, 63]:

1. Exactly 4 spikes were induced as a response to current injection of 0.696 nA for the duration of 150 ms (square pulse),
2. exactly 1 spike was induced as a response to a distal (distance from soma = 620 *μ*m) synaptic conductance (alpha-shaped with a time constant of 5 ms and a maximum amplitude of 0.0612 *μ*S),
3. the neuron responded with exactly 2 spikes to a combined stimulus of a somatic current injection (1.137 nA, 10 ms, square pulse) and a distal synaptic conductance (alpha-shaped with a time constant of 5 ms and a maximum amplitude of 0.100 *μ*S, applied 5 ms after the somatic pulse),
4. the integrated difference between the f-I curves of the variant model neuron and the control model neuron did not exceed 10% of the integral of the control neuron f-I curve, and
5. the membrane-potential limit cycle should not be too different from the control neuron limit cycle (*d_cc_*(*lc*1*, lc*2) ≤ 600, see [60] for the definition of the metric *d_cc_*).

Here, the first three conditions constrain the magnitudes of transient responses of the neuron model and the amplitudes were chosen to guarantee the largest stability with respect to the response of the model using default parameters (stable here, means that an equal change in current amplitude on a logarithmic scale is needed to produce one action potential more or one action potential less.

In the case of a violation of one or more conditions, the effect of the variant on the parameters of the model was down-scaled to a fraction *c* < 1, where the violation is observed for the first time. The fraction for each considered variant can be found in Supplementary table S2. If however an unscaled variant did not violate the conditions 1–5, we explored threshold effects up to twice the original effect, i.e., *c* ≤ 2.

Since the parameters of the neuron model subject to change by the variants were very diverse, i.e. had various different roles and dimensions (mV, mM, ms, etc.), we adopted a careful scaling strategy. Parameters that could take positive and negative values were scaled linearly with the scaling factor *c*. Parameters that were exclusively non-negative, on the other hand, were scaled logarithmically with *c*. In detail, this meant that the difference in offset potentials (Voffm, Voffh) between the control model neuron and the variant model neuron, which are described by an additive term (i.e. ±x mV), was multiplied by the down-scaling parameter c. However, the change in all other parameters, which were described by multiplicative terms, was exponentiated by the down-scaling factor *c*. This resulted in a continuous change of parameters for *c* ∈ [0, 1], which is also directly extendable to down-scaling factors > 1.

**Table 1:**
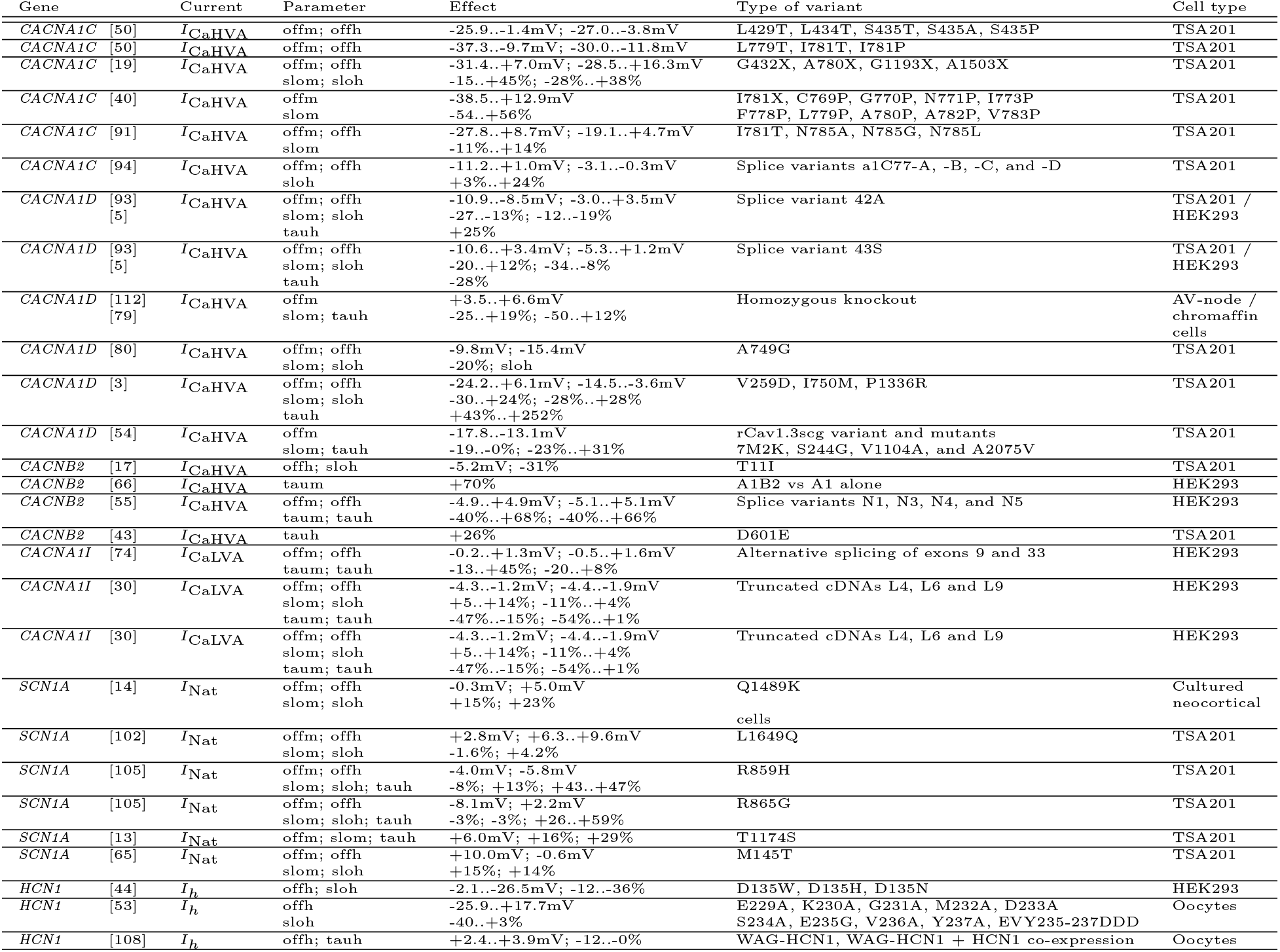
Overview over the genetic variants. The first column of the table shows the gene whose variant was studied in the named reference. Columns two and three show the current species and the affected model parameters, *offm* and *offh* representing the mid-points of activation and inactivation, respectively, *slom* and *sloh* their respective slopes, and *taum* and *tauh* their respective time constants. Multiple parameter changes in a single row are separated by a semicolon. The parameter names may refer to multiple model parameters: For example, for HVA Ca^2+^ currents, a change in *offm* means a concurrent change in parameters *V*_offma_ and *V*_offmb_, while *offh* refers to parameters *V*_offha_ and *V*_offhb_, *slom* to *V*_sloma_ and *V*_slomb_, *sloh* to *V*_sloha_ and *V*_slohb_, *taum* to *τ*_ma_ and *τ*_mb_, and *tauh* to *τ*_ha_ and *τ*_hb_. See model equations in Supplementary material for further details. The fourth column shows both direction and magnitude of the effect, where ±x mV refers to a change of the mid-point of the (in)activation curve by an absolute number, and ±x% refers to a percentual change in the underlying quantity. In many cases, studies considered more than one variant and here, they are categorized by the type of the variant where necessary (e.g., in [50] several variants of four loci, of which three were in pore-lining IS6 segment and two in bundle-crossing region of segment IIS6, were considered, and the variants are here categorized according to the segment they acted on). Column five gives the type of variant used, while the last column names the cell type under consideration.

**Table 2:**
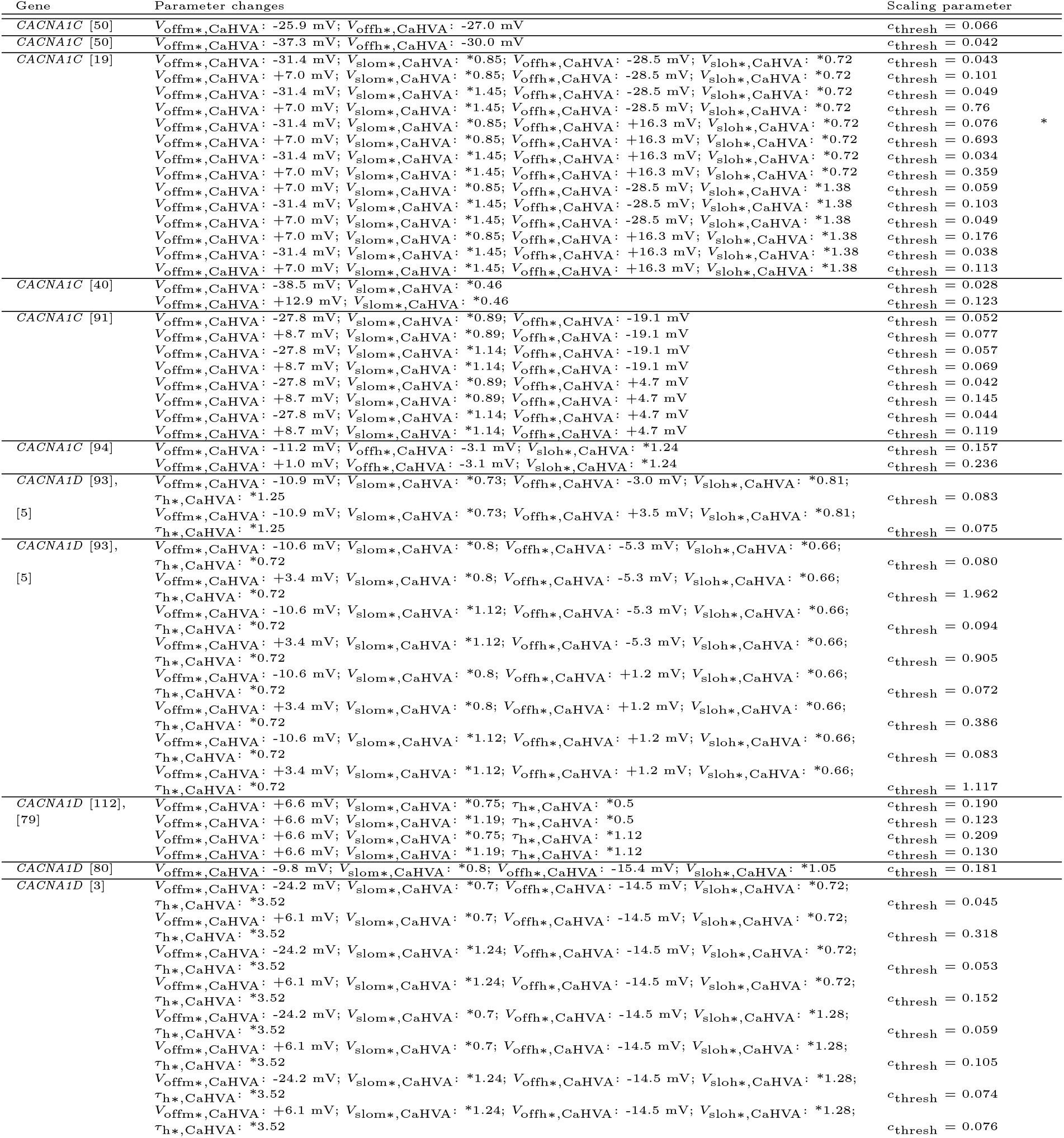
The effects of the genetic variants on model parameters. The first column names the gene in which the gene variant was analyzed and the study where the effects were reported. The second column shows the effect of the variant on the model parameters, here *offm* and *offh* represent the midpoints of activation and inactivation, respectively, *slom* and *sloh* their respective slopes, and *taum* and *tauh* their respective time constants. The third column shows the down-scaling parameter resulting from the scaling procedure outlined above. The horizontal separation of rows refers to the different entries of Supplementary table 1: If the corresponding study showed a large range of effects on single model parameters, the endpoints of such ranges are here treated as separate variants and are therefore downscaled independently of each other. The variants shown in Figures 3, 4 and 5 are marked with an asterisk.

**Table 3:**
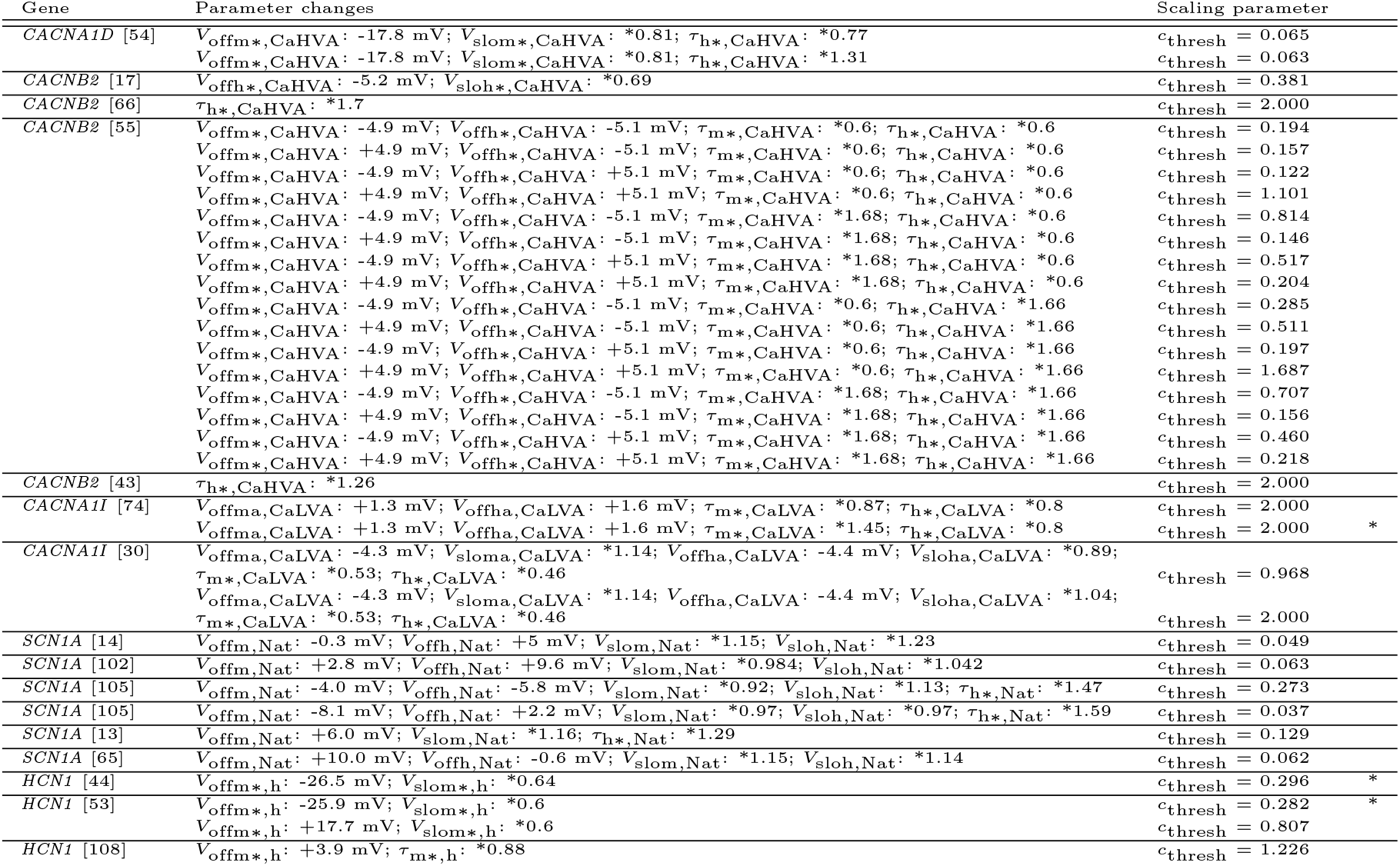

#### Synaptic Alterations

ASSR deficits in schizophrenia have previously been attributed to changes in synaptic transmission, mainly at GABAergic synapses [32]. To compare the effects of the cell-intrinsic changes to excitability introduced by the genetic variants considered in this study to these alterations at the synaptic level, we implemented the two most commonly modelled synaptic alterations [104, 70]: 1) a reduction of GABA levels due to a decrease in the expression of GAD67 [32], a GABA precursor, operationalized as a 25% reduction of the maximal GABA conductance *g_max_* at GABAergic synapses (referred to as the *Gmax* condition) and 2) an increase of the decay time constant *τ_decay_* at GABAergic synapses, as a result of a decrease of GAT1 [32], an enzyme responsible for GABA reuptake, operationalized as an increase from 8 ms to 25 ms (referred to as the *IPSC* condition).

### Data analysis

The simulated local field potential (LFP) was recorded using NetPyNE’s LFP recording capabilities with a recording time step of 0.1 ms. From the simulated LFP signals we calculated two measures capturing the degree of synchronous network activity: 1) the power spectral density and 2) the inter-trial phase coherence of the entrained network oscillation.

To compute the power spectral density, we used Welch’s periodogram method (using the implementation in matplotlib.pyplot). Specifically, the power spectral density *P_xx_* of the signal is computed as

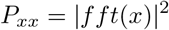

where *x* is the input signal and *fft* the Fast-Fourier Transform.

For the inter-trial phase coherence, we band-pass filtered the simulated LFP signal around the frequency of interest, i.e. 40 Hz ±2.5 Hz, and computed an analytic signal using the Hilbert transform. The analytic signal *x_a_*(*t*) can be expressed in polar coordinates as

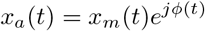

where *x_m_*(*t*) is called the instantaneous amplitude or envelope and *ϕ*(*t*) is called the instantaneous phase. We calculated the instantaneous phase *ϕ_k_*(*t*) for each trial *k* of each subject *j* and then, for each subject *j*, the inter-trial phase coherence was calculated as

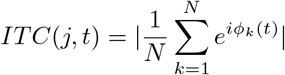

We then averaged the individual inter-trial phase coherences over time. The ITC is a normalized measure for which a value of 0 reflects maximal variability of phases across trials and a value of 1 no variability.

To further understand the effect the changes to individual parameters have on the gamma power, we correlated (Pearson correlation) the parameter changes (i.e. changes to *offm*, *offh*, *slom*, *sloh*, *taum* and *tauh*), for the model variants affecting the HVA Ca^2+^ channel with the change in evoked gamma power. Since we only included relatively few model variants affecting the other channels (LVA Ca^2+^, Nat and h), we omitted such an analysis for those channels.

Comparison of single model variants, combinations of model variants and synaptic alterations against the shared control was performed using Cumming estimation plots (see Figures 3, 4, and 5) and permutation t-tests (see Table 4) using the DABEST Python package [39]. 5000 bootstrap samples were taken; the confidence interval was bias-corrected and accelerated. The *p* values reported are the likelihoods of observing the mean differences, if the null hypothesis of zero difference is true. For each permutation *p* value, 5000 reshuffles of the control and test labels were performed. For more details see [39].

To compare our findings with our earlier findings [63], we calculated Pearson correlation coefficients between gamma power and delta resonance and PPI threshold, respectively. Specifically, delta resonance power from our earlier study was calculated as the median of the power spectrum amplitude at a baseline frequency of 1.5 Hz. The PPI threshold is the threshold synaptic conductance of 3000 simultaneously activated synapses for generating a second action potential if a suprathreshold stimulus was given to the same synapses 60 ms before. For further details see [63]. We have to note that delta resonance power in our earlier work was generated in a substantially different network model, although based on the same single cell model of layer 5 pyramidal cells, and that the PPI thresholds were generated with the full layer 5 pyramidal cell model from Hay et al. [37]. Importantly, the modelling of the SNP-like genetic variants followed exactly the same procedure as outlined here.

## Supplementary Results

### Single Cell Behaviour

The response of the two single cell models in response to somatic, and in the case of the layer 5 pyramidal cell model, also to dendritic input currents has been extensively studied in earlier studies [58, 103].

#### Network Behaviour

##### Noise-driven network

To validate our control network model, we first performed simulations (20 subjects, 10 trials each subject, see Supplementary Section 4) with only the background noise as input and calculated the background firing rates for each population. The pyramidal cell population fired at an average rate of 7.81 Hz (standard deviation: 0.07 Hz) and the basket cell population at 6.18 Hz (standard deviation: Hz), which is reasonably close to the range of 4.9 Hz (standard deviation: 0.5) reported for auditory cortex [42].

In order to explore the behaviour of the control network further, simulations with 20, 30 and 40 Hz drive were performed and the simulated local field potential was recorded. We explicitly went beyond purely driving the network with 40 Hz stimuli and included drive at 20 and 30 Hz, which are routinely used in experimental ASSR studies in patients with schizophrenia (e.g. [51, 104]). As can be seen in Supplementary Figure 7, the network clearly entrains to the driving frequency, the power at drive frequency is highest for 40 Hz and lowest for 20 Hz and for 20 Hz drive, a clear 40 Hz component is visible, thus, the network replicates experimental studies [51, 104] and previous modeling studies [104, 70, 68, 86].

**Figure 7:**
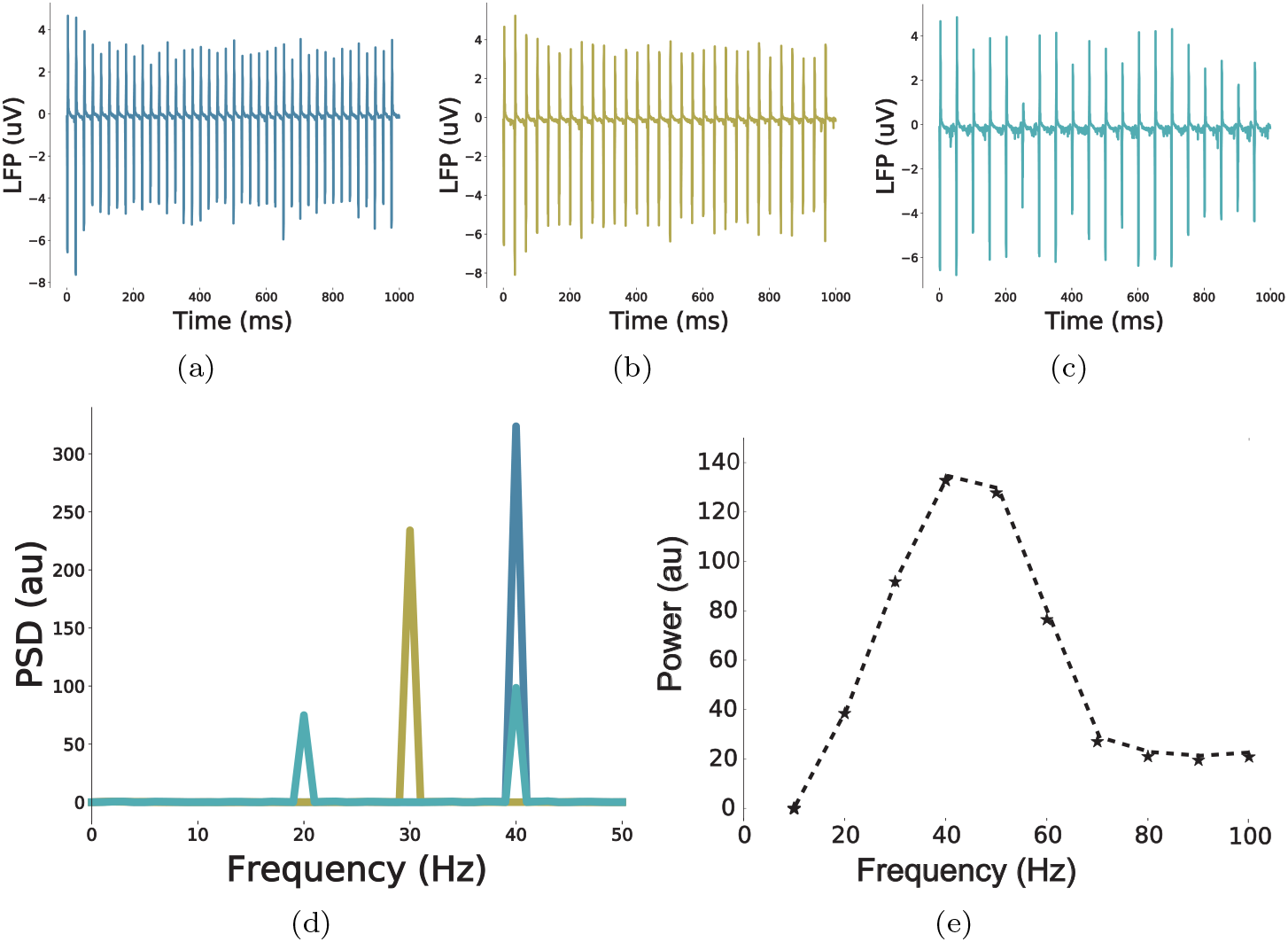
Validation of the control network model. (a) Simulated LFP in response to a 40 Hz drive. (b) Simulated LFP in response to a 30 Hz drive. (a) Simulated LFP in response to a 20 Hz drive. (d) Power spectral densities of the three signals from (a)-(c). (e) Peak power for simulations with 10-100 Hz drive to only the FS basket cell populations (Note that in the standard ASSR condition (a-d) both cell populations receive drive input).

Furthermore, Cardin et al. [11] show that rhythmic optogenetic drive of FS cells specifically enhances power in the gamma range. In our control network model, we rhythmically drive FS cells (with a sinusoidal current instead of optogenetic drive though) with frequencies between 10 and 100 Hz, in steps of 10 Hz, while excitatory cells only receive noise drive. Similarly to Cardin et al. [11], we see that periodically stimulating FS cells enhances LFP power especially in the gamma frequency range (30-60 Hz) (Supplementary Figure 7). This replicates experimental [11] and modelling findings [103] and further underpins the validity of the employed network model.

#### Single Variant Effects

Supplementary Table 4 shows the mean differences and permutation statistics for selected single model variants and model variant combinations as well as the two synaptic alterations.

**Table 4:** Permutation test statistics. Comparison against a shared control with a sample size of *n* = 20. 5000 bootstrap samples were taken; the confidence interval is bias-corrected and accelerated. The *p* values reported are the likelihoods of observing the mean differences, if the null hypothesis of zero difference is true. For each permutation *p* value, 5000 reshuffles of the control and test labels were performed. For more details see [39].

|  | sample size | mean difference | 99% CI interval | permutation p-value |
| --- | --- | --- | --- | --- |
| Ca7 | 20 | -29.5 | [-30.9, -28.2] | < 0.001 |
| HCN2 | 20 | -21.9 | [-23.7, -20.1] | < 0.001 |
| Ca74 | 20 | -16.7 | [-18.1, -15.2] | < 0.01 |
| C1 | 20 | -49.7 | [-51.0, -48.4] | < 0.001 |
| C2 | 20 | -61.9 | [-62.2, -60.6] | < 0.001 |
| C3 | 20 | -67.2 | [-68.5, -65.9] | < 0.001 |
| Reduced $g_{max}$ | 20 | -88.4 | [-90.0, -86.7] | < 0.001 |
| Increased $\tau_{inh}$ | 20 | -181.0 | [-183.0, -17.0] | < 0.001 |

Several experimental [51, 104] and modeling [104, 70, 68, 71] studies have found an increase in the subharmonic 20 Hz component in response to 40 Hz drive in SCZ. Therefore, we also quantified 20 Hz power in our exploration of the single model variant effects. As can be seen in Supplementary Figure 8, single model variants had hardly any effects on the 20 Hz component.

**Figure 8:**
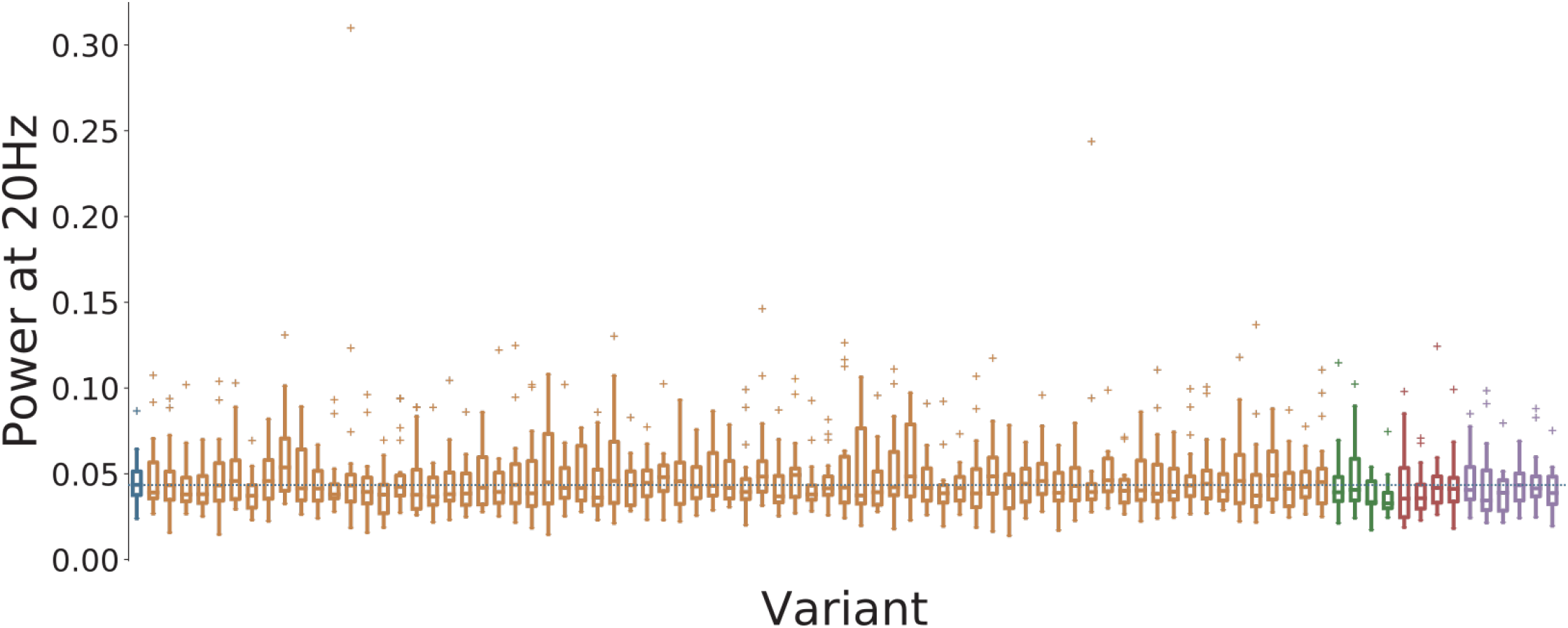
Overview of the 2040 measure for all variants. Control network in blue, *Ca*^2+^ channel variants affecting *Ca*_*HV A*_ in orange, affecting *Ca_LV A_* in green, HCN variants in red, and SCN variants in purple. Solid lines represent the mean, box edges the 25 and 75 percentile, respectively, the whisker extend to 2 standard deviations and + depict outliers. The dashed blue line represents the control mean.

**Figure 9:**
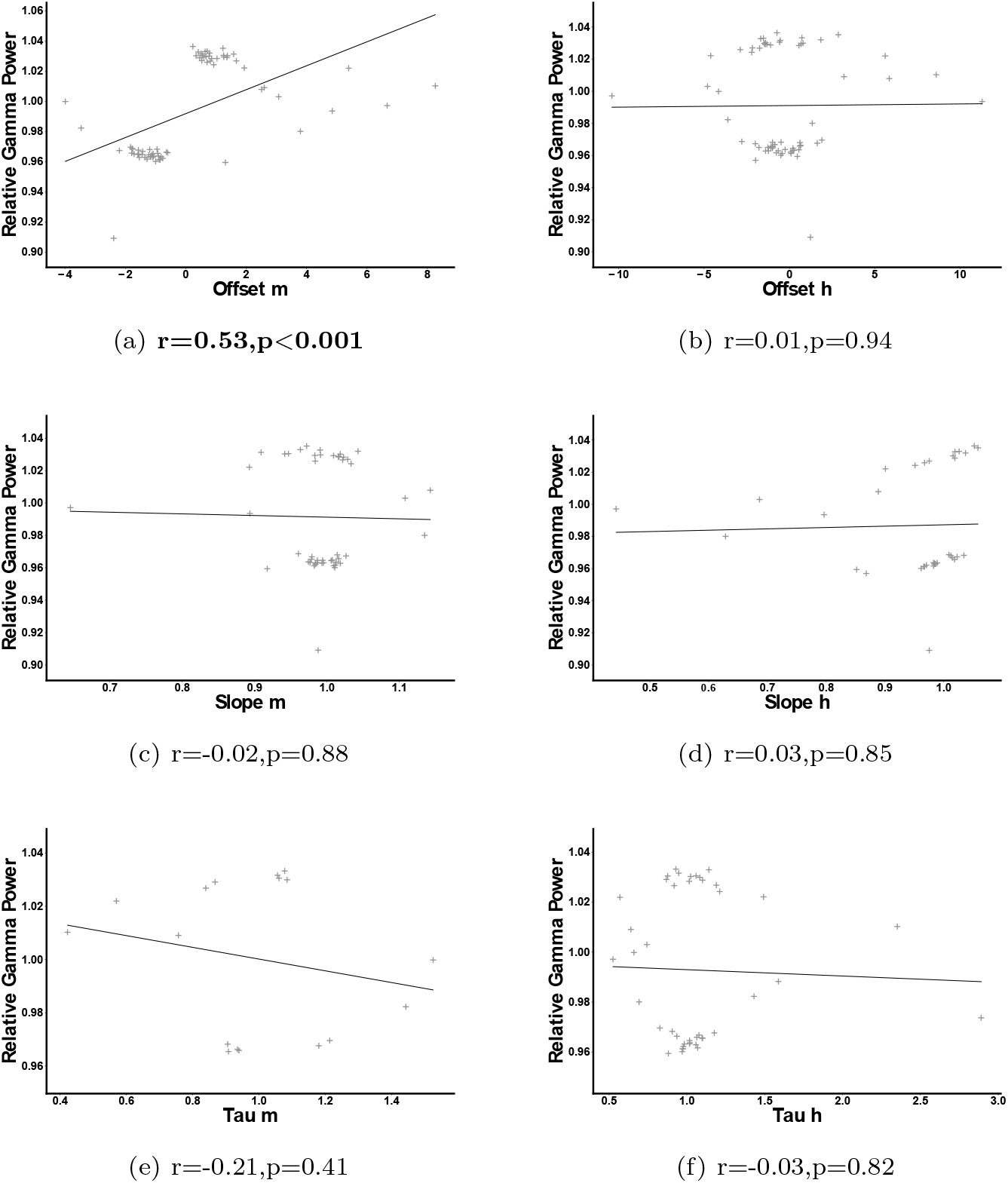
Correlation of parameter changes with gamma reduction. Pearson correlation between scaled changes of a parameter of the high-voltage activated Ca^2+^ channel and relative gamma power (i.e. the ratio of gamma power of a variant and the gamma power of the control).

Lastly, we correlated parameter changes to gamma reduction for the model variants affecting *I*_*CaHV A*_. As can be seen in Supplementary Figure S9 the change in evoked gamma power was positively correlated with the offset of the activation variable *m* but there was no significant correlation with other parameter changes.

## Data Availability

All code to simulate the computational model, to analyze the data and to create the figures is available at https://github.com/ChristophMetzner/ACnet. The model code will also be made available through ModelDB (https://senselab.med.yale.edu/modeldb/) upon publication. Furthermore, the model will also be included in the ASSRUnit package, which is designed for automated validation and comparison of models of ASSR deficits in psychiatric disorders [69].

## Notes

### Competing Interest Statement

The authors have declared no competing interest.

## References

[1] Mara Almog and Alon Korngreen. A quantitative description of dendritic conductances and its application to dendritic excitation in layer 5 pyramidal neurons. Journal of Neuroscience, 34(1):182–196, 2014.

[2] A Andrade, J Hope, A Allen, V Yorgan, D Lipscombe, and JQ Pan. A rare schizophrenia risk variant of cacna1i disrupts cav3. 3 channel activity. Scientific reports, 6:34233, 2016.

[3] Elena AB Azizan, Hanne Poulsen, Petronel Tuluc, Junhua Zhou, Michael V Clausen, Andreas Lieb, Carmela Maniero, Sumedha Garg, Elena G Bochukova, Wanfeng Zhao, et al. Somatic mutations in atp1a1 and cacna1d underlie a common subtype of adrenal hypertension. Nature Genetics, 45(9):1055–1060, 2013.

[4] Tom Binzegger, Rodney J Douglas, and Kevan AC Martin. A quantitative map of the circuit of cat primary visual cortex. Journal of Neuroscience, 24(39):8441–8453, 2004.

[5] Gabriella Bock, Mathias Gebhart, Anja Scharinger, Wanchana Jangsangthong, Perrine Busquet, Chiara Poggiani, Simone Sartori, Matteo E Mangoni, Martina J Sinnegger-Brauns, Stefan Herzig, et al. Functional properties of a newly identified c-terminal splice variant of cav1. 3 l-type ca2+ channels. Journal of Biological Chemistry, 286(49):42736–42748, 2011.

[6] Christoph Börgers and Nancy Kopell. Synchronization in networks of excitatory and inhibitory neurons with sparse, random connectivity. Neural computation, 15(3):509–538, 2003.

[7] David L Braff, Tiffany A Greenwood, Neal R Swerdlow, Gregory A Light, and Nicholas J Schork. Advances in endophenotyping schizophrenia. World Psychiatry, 7(1):11–18, 2008.

[8] Gyorgy Buzsaki. Rhythms of the Brain. Oxford University Press, 2006.

[9] György Buzsáki and Andreas Draguhn. Neuronal oscillations in cortical networks. science, 304(5679):1926–1929, 2004.

[10] Caglar Cakan and Klaus Obermayer. Biophysically grounded mean-field models of neural populations under electrical stimulation. PLoS computational biology, 16(4):e1007822, 2020.

[11] Jessica A Cardin, Marie Carlén, Konstantinos Meletis, Ulf Knoblich, Feng Zhang, Karl Deisseroth, Li-Huei Tsai, and Christopher I Moore. Driving fast-spiking cells induces gamma rhythm and controls sensory responses. Nature, 459(7247):663–667, 2009.

[12] William A Catterall, Edward Perez-Reyes, Terrance P Snutch, and Joerg Striessnig. International union of pharmacology. xlviii. nomenclature and structure-function relationships of voltage-gated calcium channels. Pharmacological Reviews, 57(4):411–425, 2005.

[13] Sandrine Cestèle, Angelo Labate, Raffaella Rusconi, Patrizia Tarantino, Laura Mumoli, Silvana Franceschetti, Grazia Annesi, Massimo Mantegazza, and Antonio Gambardella. Divergent effects of the t1174s scn1a mutation associated with seizures and hemiplegic migraine. Epilepsia, 54(5):927–935, 2013.

[14] Sandrine Cestèle, Paolo Scalmani, Raffaella Rusconi, Benedetta Terragni, Silvana Franceschetti, and Massimo Mantegazza. Self-limited hyperexcitability: Functional effect of a familial hemiplegic migraine mutation of the Nav1.1 (SCN1A) na+ channel. Journal of Neuroscience, 28(29):7273–7283, 2008.

[15] Chi-Ming A Chen, Arielle D Stanford, Xiangling Mao, Anissa Abi-Dargham, Dikoma C Shungu, Sarah H Lisanby, Charles E Schroeder, and Lawrence S Kegeles. Gaba level, gamma oscillation, and working memory performance in schizophrenia. NeuroImage: Clinical, 4:531–539, 2014.

[16] Elodie Christophe, Nathalie Doerflinger, Daniel J Lavery, Zoltán Molnár, Serge Charpak, and Etienne Audinat. Two populations of layer v pyramidal cells of the mouse neocortex: development and sensitivity to anesthetics. Journal of Neurophysiology, 94(5):3357–3367, 2005.

[17] Jonathan M Cordeiro, Mark Marieb, Ryan Pfeiffer, Kirstine Calloe, Elena Burashnikov, and Charles Antzelevitch. Accelerated inactivation of the L-type calcium current due to a mutation in CACNB2b underlies brugada syndrome. Journal of Molecular and Cellular Cardiology, 46(5):695–703, 2009.

[18] Bruce N Cuthbert and Thomas R Insel. Toward the future of psychiatric diagnosis: the seven pillars of rdoc. BMC medicine, 11(1):126, 2013.

[19] Katrin Depil, Stanislav Beyl, Anna Stary-Weinzinger, Annette Hohaus, Eugen Timin, and Steffen Hering. Timothy mutation disrupts the link between activation and inactivation in cav1. 2 protein. Journal of Biological Chemistry, 286(36):31557–31564, 2011.

[20] Alain Destexhe, Zachary F Mainen, and Terrence J Sejnowski. An efficient method for computing synaptic conductances based on a kinetic model of receptor binding. Neural computation, 6(1):14–18, 1994.

[21] Alain Destexhe, Zachary F Mainen, and Terrence J Sejnowski. Synthesis of models for excitable membranes, synaptic transmission and neuromodulation using a common kinetic formalism. Journal of computational neuroscience, 1(3):195–230, 1994.

[22] A Devor, OA Andreassen, Y Wang, T Mäki-Marttunen, OB Smeland, CC Fan, AJ Schork, D Holland, WK Thompson, A Witoelar, et al. Genetic evidence for role of integration of fast and slow neurotransmission in schizophrenia. Molecular psychiatry, 22(6):792, 2017.

[23] Franc CL Donkers, Zoë A Englander, Paul HE Tiesinga, Katherine M Cleary, Hongbin Gu, and Aysenil Belger. Reduced delta power and synchrony and increased gamma power during the p3 time window in schizophrenia. Schizophrenia research, 150(1):266–268, 2013.

[24] Aranda R Duan, Carmen Varela, Yuchun Zhang, Yinghua Shen, Lealia Xiong, Matthew A Wilson, and John Lisman. Delta frequency optogenetic stimulation of the thalamic nucleus reuniens is sufficient to produce working memory deficits: relevance to schizophrenia. Biological psychiatry, 77(12):1098–1107, 2015.

[25] Salvador Dura-Bernal, Benjamin A Suter, Padraig Gleeson, Matteo Cantarelli, Adrian Quintana, Facundo Rodriguez, David J Kedziora, George L Chadderdon, Cliff C Kerr, Samuel A Neymotin, et al. Netpyne, a tool for data-driven multiscale modeling of brain circuits. Elife, 8:e44494, 2019.

[26] Mehmet Ergen, Sonja Marbach, Andreas Brand, Canan Başar-Eroğlu, and Tamer Demiralp. P3 and delta band responses in visual oddball paradigm in schizophrenia. Neuroscience letters, 440(3):304–308, 2008.

[27] Pascal Fries. Neuronal gamma-band synchronization as a fundamental process in cortical computation. Annual review of neuroscience, 32:209–224, 2009.

[28] Pascal Fries, John H Reynolds, Alan E Rorie, and Robert Desimone. Modulation of oscillatory neuronal synchronization by selective visual attention. Science, 291(5508):1560–1563, 2001.

[29] Christopher Donald Frith. The cognitive neuropsychology of schizophrenia. Psychology press, 2014.

[30] Juan Carlos Gomora, Janet Murbartián, Juan Manuel Arias, Jung-Ha Lee, and Edward Perez-Reyes. Cloning and expression of the human t-type channel cav3. 3: insights into prepulse facilitation. Biophysical journal, 83(1):229–241, 2002.

[31] Guillermo González-Burgos, Leonid S Krimer, Nadya V Povysheva, German Barrionuevo, and David A Lewis. Functional properties of fast spiking interneurons and their synaptic connections with pyramidal cells in primate dorsolateral prefrontal cortex. Journal of neurophysiology, 93(2):942–953, 2005.

[32] Guillermo Gonzalez-Burgos and David A Lewis. Gaba neurons and the mechanisms of network oscillations: implications for understanding cortical dysfunction in schizophrenia. Schizophrenia bulletin, 34(5):944–961, 2008.

[33] Guillermo Gonzalez-Burgos and David A Lewis. Nmda receptor hypofunction, parvalbumin-positive neurons, and cortical gamma oscillations in schizophrenia. Schizophrenia bulletin, 38(5):950–957, 2012.

[34] Charles M Gray, Peter König, Andreas K Engel, and Wolf Singer. Oscillatory responses in cat visual cortex exhibit inter-columnar synchronization which reflects global stimulus properties. Nature, 338(6213):334–337, 1989.

[35] Sten Grillner. Megascience efforts and the brain. Neuron, 82(6):1209–1211, 2014.

[36] Mei-Hua Hall, Grantley Taylor, Dean F Salisbury, and Deborah L Levy. Sensory gating event–related potentials and oscillations in schizophrenia patients and their unaffected relatives. Schizophrenia bulletin, 37(6):1187–1199, 2011.

[37] Etay Hay, Sean Hill, Felix Schürmann, Henry Markram, and Idan Segev. Models of neocortical layer 5b pyramidal cells capturing a wide range of dendritic and perisomatic active properties. PLoS Computational Biology, 7:e1002107, 2011.

[38] Michael L Hines and Nicholas T Carnevale. The neuron simulation environment. Neural computation, 9(6):1179–1209, 1997.

[39] Joses Ho, Tayfun Tumkaya, Sameer Aryal, Hyungwon Choi, and Adam Claridge-Chang. Moving beyond p values: Everyday data analysis with estimation plots. Nature Methods, 16:565–566, 2019.

[40] Annette Hohaus, Stanislav Beyl, Michaela Kudrnac, Stanislav Berjukow, Eugen N Timin, Rainer Marksteiner, Marion A Maw, and Steffen Hering. Structural determinants of l-type channel activation in segment iis6 revealed by a retinal disorder. Journal of Biological Chemistry, 280(46):38471–38477, 2005.

[41] L Elliot Hong, Ann Summerfelt, Braxton D Mitchell, Robert P McMahon, Ikwunga Wonodi, Robert W Buchanan, and Gunvant K Thaker. Sensory gating endophenotype based on its neural oscillatory pattern and heritability estimate. Archives of general psychiatry, 65(9):1008–1016, 2008.

[42] Tomáš Hromádka, Michael R DeWeese, and Anthony M Zador. Sparse representation of sounds in the unanesthetized auditory cortex. PLoS biology, 6(1), 2008.

[43] Dan Hu, Hector Barajas-Martinez, Vladislav V Nesterenko, Ryan Pfeiffer, Alejandra Guerchicoff, Jonathan M Cordeiro, Anne B Curtis, Guido D Pollevick, Yuesheng Wu, Elena Burashnikov, et al. Dual variation in scn5a and cacnb2b underlies the development of cardiac conduction disease without brugada syndrome. Pacing and clinical electrophysiology, 33(3):274–285, 2010.

[44] Takahiro M Ishii, Noriyuki Nakashima, and Harunori Ohmori. Tryptophan-scanning mutagenesis in the s1 domain of mammalian hcn channel reveals residues critical for voltage-gated activation. Journal of Physiology, 579(2):291–301, 2007.

[45] Kenneth S Kendler. Explanatory models for psychiatric illness. American Journal of Psychiatry, 165(6):695–702, 2008.

[46] Kübra Komek Kirli, GB Ermentrout, and Raymond Y Cho. Computational study of nmda conductance and cortical oscillations in schizophrenia. Frontiers in computational neuroscience, 8:133, 2014.

[47] Kübra Kömek, G Bard Ermentrout, Christopher P Walker, and Raymond Y Cho. Dopamine and gamma band synchrony in schizophrenia–insights from computational and empirical studies. European Journal of Neuroscience, 36(2):2146–2155, 2012.

[48] Kübra Kömek, George B Ermentrout, and Raymond Y Cho. Dopamine-nmda interactions and relevance to gamma band synchrony in schizophrenia. BMC neuroscience, 14(S1):P218, 2013.

[49] Giri P Krishnan, William P Hetrick, CA Brenner, Anantha Shekhar, AN Steffen, and Brian F O’Donnell. Steady state and induced auditory gamma deficits in schizophrenia. Neuroimage, 47(4):1711–1719, 2009.

[50] Michaela Kudrnac, Stanislav Beyl, Annette Hohaus, Anna Stary, Thomas Peterbauer, Eugen Timin, and Steffen Hering. Coupled and independent contributions of residues in IS6 and IIS6 to activation gating of CaV1.2. Journal of Biological Chemistry, 284(18):12276–12284, 2009.

[51] Jun Soo Kwon, Brian F O’Donnell, Gene V Wallenstein, Robert W Greene, Yoshio Hirayasu, Paul G Nestor, Michael E Hasselmo, Geoffrey F Potts, Martha E Shenton, and Robert W McCarley. Gamma frequency–range abnormalities to auditory stimulation in schizophrenia. Archives of general psychiatry, 56(11):1001–1005, 1999.

[52] S Hong Lee, Teresa R DeCandia, Stephan Ripke, Jian Yang, Patrick F Sullivan, Michael E Goddard, Matthew C Keller, Peter M Visscher, Naomi R Wray, Schizophrenia Psychiatric Genome-Wide Association Study Consortium, et al. Estimating the proportion of variation in susceptibility to schizophrenia captured by common snps. Nature Genetics, 44(3):247–250, 2012.

[53] Heinte Lesso and Ronald A Li. Helical secondary structure of the external s3-s4 linker of pacemaker (hcn) channels revealed by site-dependent perturbations of activation phenotype. Journal of Biological Chemistry, 278(25):22290–22297, 2003.

[54] Andreas Lieb, Anja Scharinger, Simone Sartori, Martina J Sinnegger-Brauns, and Jörg Striessnig. Structural determinants of cav1. 3 l-type calcium channel gating. Channels, 6(3):197–205, 2012.

[55] Sabine Link, Marcel Meissner, Brigitte Held, Andreas Beck, Petra Weissgerber, Marc Freichel, and Veit Flockerzi. Diversity and developmental expression of L-type calcium channel *β*2 proteins and their influence on calcium current in murine heart. Journal of Biological Chemistry, 284(44):30129–30137, 2009.

[56] Jenny Y Ma, William A Catterall, and Todd Scheuer. Persistent sodium currents through brain sodium channels induced by g protein *β γ* subunits. Neuron, 19(2):443–452, 1997.

[57] Zachary F Mainen and Terrence J Sejnowski. Influence of dendritic structure on firing pattern in model neocortical neurons. Nature, 382(6589):363–366, 1996.

[58] T Mäki-Marttunen, G Halnes, A Devor, C Metzner, A M Dale, O A Andreassen, and G T Einevoll. A stepwise neuron model fitting procedure designed for recordings with high spatial resolution: Application to layer 5 pyramidal cells. Journal of Neuroscience Methods, 273:264–283, 2018.

[59] Tuomo Mäki-Marttunen, Geir Halnes, Anna Devor, Christoph Metzner, Anders M Dale, Ole A Andreassen, and Gaute T Einevoll. A stepwise neuron model fitting procedure designed for recordings with high spatial resolution: application to layer 5 pyramidal cells. Journal of neuroscience methods, 293:264–283, 2018.

[60] Tuomo Mäki-Marttunen, Geir Halnes, Anna Devor, Aree Witoelar, Francesco Bettella, Srdjan Djurovic, Yunpeng Wang, Gaute T. Einevoll, Ole A. Andreassen, and Anders M. Dale. Functional effects of schizophrenia-linked genetic variants on intrinsic single-neuron excitability: A modeling study. Biological Psychiatry: Cognitive Neuroscience and Neuroimaging, 1:49–59, 2016.

[61] Tuomo Mäki-Marttunen, Nicolangelo Iannella, Andrew G Edwards, Gaute Einevoll, and Kim T Blackwell. A unified computational model for cortical post-synaptic plasticity. eLife, 2020.

[62] Tuomo Mäki-Marttunen, Tobias Kaufmann, Torbjørn Elvsåshagen, Anna Devor, Srdjan Djurovic, Lars T Westlye, Marja-Leena Linne, Marcella Rietschel, Dirk Schubert, Stefan Borgwardt, et al. Biophysical psychiatry—how computational neuroscience can help to understand the complex mechanisms of mental disorders. Frontiers in psychiatry, 10, 2019.

[63] Tuomo Mäki-Marttunen, Florian Krull, Francesco Bettella, Espen Hagen, Solveig Næss, Torbjørn V Ness, Torgeir Moberget, Torbjørn Elvsåshagen, Christoph Metzner, Anna Devor, et al. Alterations in schizophrenia-associated genes can lead to increased power in delta oscillations. Cerebral Cortex, 29(2):875–891, 2019.

[64] Tuomo Mäki-Marttunen, Glenn T. Lines, Andrew G. Edwards, Aslak Tveito, Anders M. Dale, Gaute T. Einevoll, and Ole A. Andreassen. Pleiotropic effects of schizophrenia-associated genetic variants in neuron firing and cardiac pacemaking revealed by computational modeling. Translational Psychiatry, 7:5, 2017.

[65] Massimo Mantegazza, Antonio Gambardella, Raffaella Rusconi, Emanuele Schiavon, Ferdinanda Annesi, Rita Restano Cassulini, Angelo Labate, Sara Carrideo, Rosanna Chifari, Maria Paola Canevini, et al. Identification of an nav1.1 sodium channel (scn1a) loss-of-function mutation associated with familial simple febrile seizures. Proceedings of the National Academy of Sciences of the United States of America, 102(50):18177–18182, 2005.

[66] Enrique Massa, Kevin M Kelly, David I Yule, Robert L MacDonald, and Michael D Uhler. Comparison of fura-2 imaging and electrophysiological analysis of murine calcium channel alpha 1 subunits coexpressed with novel beta 2 subunit isoforms. Molecular Pharmacology, 47(4):707–716, 1995.

[67] Lucia Melloni, Carlos Molina, Marcela Pena, David Torres, Wolf Singer, and Eugenio Rodriguez. Synchronization of neural activity across cortical areas correlates with conscious perception. Journal of neuroscience, 27(11):2858–2865, 2007.

[68] Christoph Metzner. [Re] Modeling GABA Alterations in Schizophrenia: A Link Between Impaired Inhibition and Gamma and Beta Auditory Entrainment. ReScience, 3(1):6, August 2017.

[69] Christoph Metzner, Tuomo Mäki-Marttunen, Bartosz Zurowski, and Volker Steuber. Modules for automated validation and comparison of models of neurophysiological and neurocognitive biomarkers of psychiatric disorders: Assrunit—a case study. Computational Psychiatry, 2:74–91, 2018.

[70] Christoph Metzner, Achim Schweikard, and Bartosz Zurowski. Multifactorial modeling of impairment of evoked gamma range oscillations in schizophrenia. Frontiers in computational neuroscience, 10:89, 2016.

[71] Christoph Metzner, Bartosz Zurowski, and Volker Steuber. the role of parvalbumin-positive interneurons in auditory steady-state response deficits in schizophrenia. Scientific Reports, 9(1):1–16, 2019.

[72] Patricia T Michie, Hamish Innes-Brown, Juanita Todd, and Assen V Jablensky. Duration mismatch negativity in biological relatives of patients with schizophrenia spectrum disorders. Biological psychiatry, 52(7):749–758, 2002.

[73] A Munoz, Timothy M Woods, and Edward G Jones. Laminar and cellular distribution of ampa, kainate, and nmda receptor subunits in monkey sensory– motor cortex. Journal of Comparative Neurology, 407(4):472–490, 1999.

[74] Janet Murbartián, Juan Manuel Arias, and Edward Perez-Reyes. Functional impact of alternative splicing of human t-type cav3. 3 calcium channels. Journal of neurophysiology, 92(6):3399–3407, 2004.

[75] Chaelon IO Myme, Ken Sugino, Gina G Turrigiano, and Sacha B Nelson. The nmda-to-ampa ratio at synapses onto layer 2/3 pyramidal neurons is conserved across prefrontal and visual cortices. Journal of neurophysiology, 90(2):771–779, 2003.

[76] Athanasia Papoutsi, Kyriaki Sidiropoulou, Vassilis Cutsuridis, and Panayiota Poirazi. Induction and modulation of persistent activity in a layer v pfc micro-circuit model. Frontiers in neural circuits, 7:161, 2013.

[77] Sohee Park and Philip S Holzman. Schizophrenics show spatial working memory deficits. Archives of general psychiatry, 49(12):975–982, 1992.

[78] Rodrigo Pavão, Adriano BL Tort, and Olavo B Amaral. Multifactoriality in psychiatric disorders: A computational study of schizophrenia. Schizophrenia bulletin, 41(4):980–988, 2015.

[79] Alberto Pérez-Alvarez, Alicia Hernández-Vivanco, Jose Carlos Caba-González, and Almudena Albillos. Different roles attributed to cav1 channel subtypes in spontaneous action potential firing and fine tuning of exocytosis in mouse chromaffin cells. Journal of Neurochemistry, 116(1):105–121, 2011.

[80] Alexandra Pinggera, Andreas Lieb, Bruno Benedetti, Michaela Lampert, Stefania Monteleone, Klaus R Liedl, Petronel Tuluc, and Jörg Striessnig. Cacna1d de novo mutations in autism spectrum disorders activate cav1. 3 l-type calcium channels. Biological psychiatry, 77(9):816–822, 2015.

[81] Astrid A Prinz, Dirk Bucher, and Eve Marder. Similar network activity from disparate circuit parameters. Nature neuroscience, 7(12):1345–1352, 2004.

[82] Davide Reato, Asif Rahman, Marom Bikson, and Lucas C Parra. Low-intensity electrical stimulation affects network dynamics by modulating population rate and spike timing. Journal of Neuroscience, 30(45):15067–15079, 2010.

[83] S. Ripke, B. M. Neale, A. Corvin, J. T. Walters, K. H. Farh, P.A. Holmans, P. Lee, B. Bulik-Sullivan, D. A. Collier, H. Huang, T.H. Pers, I. Agartz, E. Agerbo, M. Albus, M. Alexander, F. Amin, et al. Biological insights from 108 schizophrenia-associated genetic loci. Nature, 511(7510):421–427, 2014.

[84] Barbara Rosati and David McKinnon. Regulation of ion channel expression. Circulation Research, 94(7):874–883, 2004.

[85] Hyeon Seo, Natalie Schaworonkow, Sung Chan Jun, and Jochen Triesch. A multi-scale computational model of the effects of tms on motor cortex. F1000Research, 5, 2016.

[86] Peter J Siekmeier et al. Development of antipsychotic medications with novel mechanisms of action based on computational modeling of hippocampal neuropathology. PloS one, 8(3), 2013.

[87] Wolf Singer. Neuronal synchrony: a versatile code for the definition of relations? Neuron, 24(1):49–65, 1999.

[88] Nathan G Skene, Julien Bryois, Trygve E Bakken, Gerome Breen, James J Crowley, Héléna A Gaspar, Paola Giusti-Rodriguez, Rebecca D Hodge, Jeremy A Miller, Ana B Muñoz-Manchado, et al. Genetic identification of brain cell types underlying schizophrenia. Nature genetics, 50(6):825–833, 2018.

[89] Kevin M Spencer. The functional consequences of cortical circuit abnormalities on gamma oscillations in schizophrenia: insights from computational modeling. Frontiers in human neuroscience, 3:33, 2009.

[90] Kevin M Spencer, Paul G Nestor, Margaret A Niznikiewicz, Dean F Salisbury, Martha E Shenton, and Robert W McCarley. Abnormal neural synchrony in schizophrenia. Journal of Neuroscience, 23(19):7407–7411, 2003.

[91] Anna Stary, Michaela Kudrnac, Stanislav Beyl, Annette Hohaus, Eugen Timin, Peter Wolschann, H Robert Guy, and Steffen Hering. Molecular dynamics and mutational analysis of a channelopathy mutation in the iis6 helix of cav1. 2. Channels, 2(3):216–223, 2008.

[92] Catherine Tallon-Baudry, Olivier Bertrand, Franck Peronnet, and Jacques Pernier. Induced *γ*-band activity during the delay of a visual short-term memory task in humans. Journal of Neuroscience, 18(11):4244–4254, 1998.

[93] Bao Zhen Tan, Fengli Jiang, Ming Yeong Tan, Dejie Yu, Hua Huang, Yiru Shen, and Tuck Wah Soong. Functional characterization of alternative splicing in the c terminus of l-type cav1. 3 channels. Journal of Biological Chemistry, 286(49):42725–42735, 2011.

[94] Zhen Zhi Tang, Mui Cheng Liang, Songqing Lu, Dejie Yu, Chye Yun Yu, David T Yue, and Tuck Wah Soong. Transcript scanning reveals novel and extensive splice variations in human l-type voltage-gated calcium channel, cav1. 2 *α*1 subunit. Journal of Biological Chemistry, 279(43):44335–44343, 2004.

[95] Hanna Thuné, Marc Recasens, and Peter J Uhlhaas. The 40-hz auditory steady-state response in patients with schizophrenia: a meta-analysis. JAMA psychiatry, 73(11):1145–1153, 2016.

[96] B. I. Turetsky, M. E. Calkins, G. A. Light, A. Olincy, A. D. Radant, and N. R. Swerdlow. Neurophysiological endophenotypes of schizophrenia: the viability of selected candidate measures. Schizophrenia Bulletin, 33(1):69–94, 2007.

[97] Peter Uhlhaas, Gordon Pipa, Bruss Lima, Lucia Melloni, Sergio Neuenschwander, Danko Nikolić, and Wolf Singer. Neural synchrony in cortical networks: history, concept and current status. Frontiers in integrative neuroscience, 3:17, 2009.

[98] Peter J Uhlhaas and Steven M Silverstein. Perceptual organization in schizophrenia spectrum disorders: empirical research and theoretical implications. Psychological bulletin, 131(4):618, 2005.

[99] Peter J Uhlhaas and Wolf Singer. Abnormal neural oscillations and synchrony in schizophrenia. Nature reviews neuroscience, 11(2):100–113, 2010.

[100] Daniel Umbricht and Sanya Krljes. Mismatch negativity in schizophrenia: a meta-analysis. Schizophrenia research, 76(1):1–23, 2005.

[101] CFMG Van Kesteren, H Gremmels, LD De Witte, EM Hol, AR Van Gool, PG Falkai, RS Kahn, and IEC Sommer. Immune involvement in the pathogenesis of schizophrenia: a meta-analysis on postmortem brain studies. Translational psychiatry, 7(3):e1075–e1075, 2017.

[102] Kaate RJ Vanmolkot, Elena Babini, Boukje de Vries, Anine H Stam, Tobias Freilinger, Gisela M Terwindt, Lisa Norris, Joost Haan, Rune R Frants, Nabih M Ramadan, et al. The novel p.L1649Q mutation in the SCN1A epilepsy gene is associated with familial hemiplegic migraine: genetic and functional studies. Human Mutation, 28(5):522–522, 2007.

[103] Dorea Vierling-Claassen, Jessica Cardin, Christopher I Moore, and Stephanie R Jones. Computational modeling of distinct neocortical oscillations driven by cell-type selective optogenetic drive: separable resonant circuits controlled by low-threshold spiking and fast-spiking interneurons. Frontiers in human neuroscience, 4:198, 2010.

[104] Dorea Vierling-Claassen, Peter Siekmeier, Steven Stufflebeam, and Nancy Kopell. Modeling gaba alterations in schizophrenia: a link between impaired inhibition and altered gamma and beta range auditory entrainment. Journal of neurophysiology, 99(5):2656–2671, 2008.

[105] Linda Volkers, Kristopher M Kahlig, Nienke E Verbeek, Joost HG Das, Marjan JA van Kempen, Hans Stroink, Paul Augustijn, Onno van Nieuwenhuizen, Dick Lindhout, Alfred L George, et al. Nav1. 1 dysfunction in genetic epilepsy with febrile seizures-plus or dravet syndrome. European Journal of Neuroscience, 34(8):1268–1275, 2011.

[106] Vladislav Volman, M Margarita Behrens, and Terrence J Sejnowski. Downregulation of parvalbumin at cortical gaba synapses reduces network gamma oscillatory activity. Journal of Neuroscience, 31(49):18137–18148, 2011.

[107] Xiao-Jing Wang and John H Krystal. Computational psychiatry. Neuron, 84(3):638–654, 2014.

[108] Konstantin Wemhöner, Tatyana Kanyshkova, Nicole Silbernagel, Juncal Fernandez-Orth, Stefan Bittner, Aytug K Kiper, Susanne Rinné, Michael F Netter, Sven G Meuth, Thomas Budde, et al. An n-terminal deletion variant of hcn1 in the epileptic wag/rij strain modulates hcn current densities. Frontiers in molecular neuroscience, 8:83, 2015.

[109] Valérie Wespatat, Frank Tennigkeit, and Wolf Singer. Phase sensitivity of synaptic modifications in oscillating cells of rat visual cortex. Journal of Neuroscience, 24(41):9067–9075, 2004.

[110] William RJ Whitaker, Jeffrey J Clare, Andrew J Powell, Yu Hua Chen, Richard LM Faull, and Piers C Emson. Distribution of voltage-gated sodium channel *α*-subunit and *β*-subunit mrnas in human hippocampal formation, cortex, and cerebellum. Journal of Comparative Neurology, 422(1):123–139, 2000.

[111] Miles A Whittington, Mark O Cunningham, Fiona EN LeBeau, Claudia Racca, and Roger D Traub. Multiple origins of the cortical gamma rhythm. Developmental neurobiology, 71(1):92–106, 2011.

[112] Q. Zhang, V. Timofeyev, H. Qiu, L. Lu, N. Li, A. Singapuri, C. L. Torado, H. S. Shin, and N. Chiamvimonvat. Expression and roles of Cav1.3 (*α*1D) L-type Ca2+ channel in atrioventricular node automaticity. Journal of Molecular and Cellular Cardiology, 50(1):194–202, 2011.

